# Isomerization of antimalarial drug WR99210 explains its inactivity in a commercial stock

**DOI:** 10.1101/2020.07.01.182402

**Authors:** T. Parks Remcho, Sravanthi D. Guggilapu, Phillip Cruz, Glenn A. Nardone, Gavin Heffernan, Robert D. O’Connor, Carole A. Bewley, Thomas E. Wellems, Kristin D. Lane

## Abstract

WR99210, a former antimalarial drug candidate now widely used for the selection of *Plasmodium* transfectants, selectively targets the parasite dihydrofolate reductase thymidine synthase bifunctional enzyme (DHFR-TS) but not human DHFR, which is not fused with TS. Accordingly, WR99210 and plasmids expressing human *dhfr* have become valued tools for the genetic modification of parasites in the laboratory. Concerns over the ineffectiveness of WR99210 from some sources encouraged us to investigate the biological and chemical differences of supplies from two different companies (compounds **1** and **2**). Compound **1** proved effective at low nanomolar concentrations against *Plasmodium falciparum* parasites, whereas compound **2** was ineffective even at micromolar concentrations. Intact and fragmented mass spectra indicated identical molecular formulae of the unprotonated (free base) structures of **1** and **2**; however, the compounds displayed differences by thin layer chromatography, reverse phase high performance liquid chromatography, and ultraviolet-visible spectroscopy, indicating important isomeric differences. Structural evaluations by ^1^H, ^13^C, and ^15^N nuclear magnetic resonance spectroscopy confirmed **1** as WR99210 and **2** as an isomeric dihydrotriazine. Induced fit, computational docking models showed that **1** binds tightly and specifically in the *P. falciparum* DHFR active site whereas **2** fits poorly to the active site in loose and varied orientations. Stocks and concentrates of WR99210 should be monitored for the presence of isomer **2**, particularly when they are not supplied as the hydrochloride salt or are exposed to basic conditions that can promote isomerization. Absorption spectroscopy may serve for assays of the unrearranged and rearranged triazines.

## INTRODUCTION

WR99210 [4,6-diamino-1,2-dihydro-2,2-dimethyl-1-(2,4,5-trichlorophenoxypropyloxy)-1,3,5 triazine] is a folate pathway antagonist with potent activity against *Plasmodium* malaria parasites (1). Action of WR99210, originally designated as BRL 6231, in parasites is against the bifunctional dihydrofolate reductase thymidylate synthase enzyme (DHFR-TS), where the compound binds to the DHFR active site and blocks the production of tetrahydrofolate in the folate pathway (2). In contrast, WR99210 interacts only weakly with human DHFR (hDHFR). Episomal expression of the hDHFR coding sequence in *Plasmodium falciparum* produces a 4–log increase in WR99210 half-maximal effective concentration (EC_50_) against transformed parasites, providing a powerful selectable resistance marker for transfection studies (3). The exciting potential of WR99210 as an antimalarial drug candidate eventually waned after preclinical and clinical trials demonstrated adverse events including severe gastrointestinal side effects in non-human primates and humans, even at low drug doses (4-7). Pursuit of a pro-drug form of WR99210 (PS-15) was also limited by a regulatory restriction on the starting material, 2,4,5 trichlorophenol, used to produce dioxin, among other toxic substances (8, 9). However, alternative antifolate compounds that incorporate a flexible linker as seen in WR99210 are under evaluation, including candidate P218 which is in clinical trials with the Medicines for Malaria Venture (10-13).

Genetic transfections and transformations of *Plasmodium* have become elemental tools of malaria research. Positive selection of hDHFR-transformed parasites is commonly achieved by WR99210 exposure *in vitro* and *in vivo* with *P. falciparum* as well as certain other *Plasmodium* species (14-16). Untransformed *Plasmodium spp.* are sensitive to nanomolar levels of WR99210, and spontaneous development of WR99210 resistance has not been reported from exposure to the compound (17). The usefulness of the compound for the selection of hDHFR-transformed lines thus created an ongoing demand for WR99210 for genetic manipulations of the parasites.

In recent transfection experiments, we observed that WR99210 from one source (stock JP, from Jacobus Pharmaceutical Company Inc., Princeton, NJ) was effective at nanomolar concentrations, but a stock of WR99210 from another source (stock SA, from Sigma-Aldrich Corp. – St Louis, USA) did not kill untransformed *P. falciparum* parasites, even at micromolar concentrations. After this discovery, and learning of similar observations reported in online scientific discussion groups (18, 19), we initiated investigations into potential differences between the JP and SA WR99210 stock compounds. Here we report results from drug response assays, chemical and structural evaluations, and computational modeling studies that demonstrate dramatic effects on activity from isomeric differences between the JP and SA stocks. Further, we propose a mechanism by which the inactive isomer may develop from the active WR99210 compound, and we suggest some methods for rapid detection of the inactive isomer.

## RESULTS

### *P. falciparum* parasites exhibit strikingly different responses to stocks of WR99210 from two sources

For more than 20 years (3), our standard EC_50_ assays with JP WR99210 have routinely yielded susceptibilities in the sub-nanomolar range for untransformed *P. falciparum* parasites. Results for two *P. falciparum* lines are listed in Table 1. NF54 encodes an antifolate sensitive isoform of PfDHFR (N51, C59, S108), while Dd2 (N51I, C59R, S108N) encodes mutations at PfDHFR residues that confer resistance to certain other antifolates such as pyrimethamine, but not to WR99210. NF54 EC_50_ results with JP WR99210 were 0.056 nM with a 95% confidence interval (CI) of 0.029 – 0.103 nM, while Dd2 showed a 10× increase to 0.62 nM (CI: 0.580 – 0.671 nM). In striking contrast to these results with JP WR99210, our EC_50_ results from stocks of more recently acquired SA WR99210 (Sigma catalog no. 47326-86-3 lots/batches 0000042122, 0000014239, 095M4622V, 070M4610V) were 900 – 23,000 fold higher: 1.25 µM (CI: 1.01 – 1.57 µM) for NF54 and 547 nM (CI: 525 – 571 nM) for Dd2 (Table 1).

**Table 1.** WR99210 efficacy differs based on source

| Strain | Source | EC<sub>50</sub> | n | Fold Change | PfDHFR Genotype |  |  |
| --- | --- | --- | --- | --- | --- | --- | --- |
|  |  | 95% CI |  |  | 51 | 59 | 108 |
| NF54 | JP | 0.029 to 0.103 nM | 2 |  | N | C | S |
|  | SA | 1.01 to 1.57 µM | 4 | 22,500 |  |  |  |
| Dd2 | JP | 0.580 to 0.671 nM | 4 |  | I | R | N |
|  | SA | 525 to 571 nM | 4 | 887 |  |  |  |

JP: Jacobus Pharmaceutical, SA: Sigma Aldrich, SD: Standard deviation, Fold Change = SA EC<sub>50</sub>/JP EC<sub>50</sub>

Morphological appearances of parasites exposed to stock compounds of JP WR99210 and SA WR99210 were consistent with results of the EC_50_ assays. Culture samples of synchronized NF54 schizont-infected erythroctyes were exposed to 2 nM WR99210 from JP or SA, to 1 µM WR99210 from SA, or to 0.0002% DMSO (control) for 4 days. Microscopy of Giemsa-stained thin smears of these exposed cells showed dead and dying parasites only in response to 2 nM JP WR99210; the appearances of the parasitized cells under all of the other exposures remained healthy, including parasites exposed to 1 µM SA WR99210.

### WR99210 stocks from the two sources show no discernable differences by high resolution mass spectrometry

To determine whether the JP and SA stocks of WR99210 differed in their chemical composition, we compared dimethyl sulfoxide (DMSO)-solubilized samples of each stock by high resolution mass spectrometry (HRMS).The HRMS spectra for JP WR99210 (compound **1**), supplied as the hydrochloride (HCl) salt, and SA WR99210 (compound **2**), supplied as free base, were similar, showing quasi-molecular ions, of atomic mass units 394.0604 and 394.0606, respectively, both consistent with the WR99210 molecular formula of C_14_H_18_Cl_3_N_5_O_2_ (Fig. 1A; Fig. S1-2). HRMS analysis of the concentrates used for the drug response assays confirmed that both compounds were present in the expected levels.

**Figure 1.**
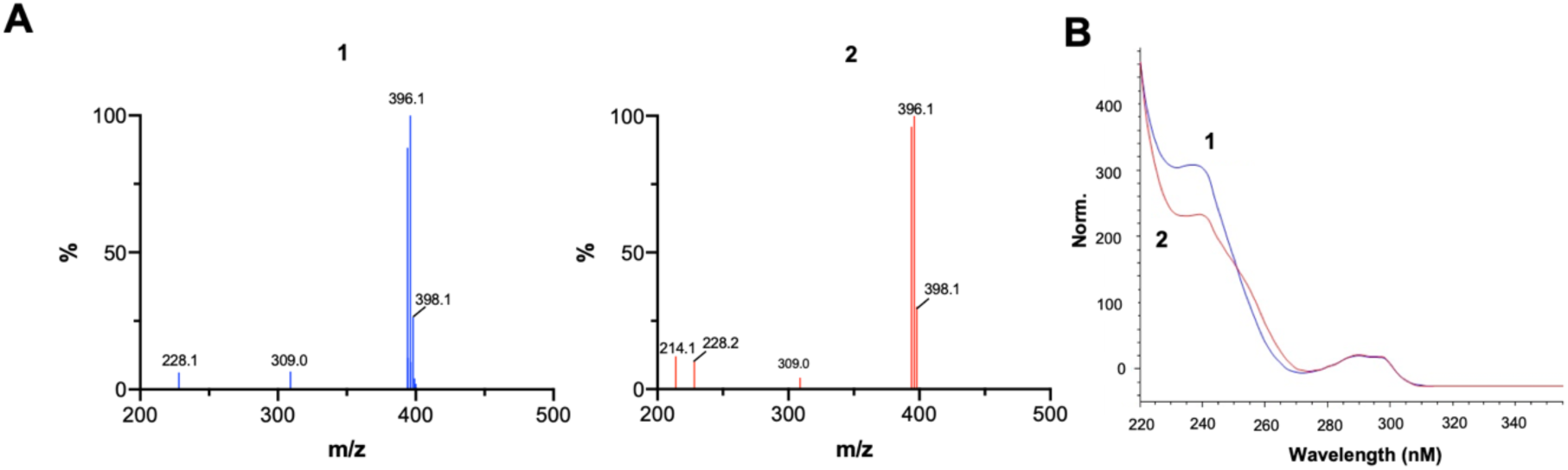
WR99210 (1) and the inactive isomer (2) cannot be discriminated by HRMS but isomeric differences are observed by UV-vis spectroscopy. (A) HRMS shows a lack of substantial contaminants or degrative products and shows equivalent quasi-molecular ions. (B) Discrimination of **1** and **2** by spectroscopy at ultraviolet wavelengths. The 230 – 240 nm plateau and shoulder region between 250 and 260 nm indicated the presence of active WR99210 (**1**) and identified the presence of the inactive isomer (**2**).

### An isomeric difference is suggested by TLC, RP-HPLC, and UV-vis spectroscopy

We next checked for evidence of a structural difference between **1** and **2** that could explain their different effects in the drug response assays. Thin-layer chromotography (TLC), reverse phase high performance liquid chromatography (RP-HPLC), and ultraviolet-visible spectroscopy (UV-vis) were performed with the samples dissolved in tetrahydrofuran:methanol (2:1), which provided solublities of more than 3 mg/mL *vs.* 0.2 mg/ml with DMSO. Silica gel TLC of these samples developed with a chloroform:diethyl ether:methanol solvent system showed different retention factors (*R*_f_) for **1** (*R*_f_ = 0.30) and **2** (*R*_f_ = 0.37) (Fig. S3). RP-HPLC experiments indicated that both compounds were more than 95% pure and eluted with different retention times (*t*_R_) of 13.7 minutes for **1** and 13.1 minutes for **2** (Fig. S4-5).

Consistent with the differences between **1** and **2** detected by TLC and RP-HPLC, UV-vis spectroscopy showed the presence of two separate compounds. These differences were particularly evident in the different absorbances in the plateau region between 230 – 240 nm (Fig. 1B). The results of these analyses and the drug assays pointed to the likelihood that **2** was an inactive isomer of the WR99210 structure **1**.

### Chemical derivatization and nuclear magnetic resonance spectroscopy clarify structural differences between compounds 1 and 2

We next sought to determine the structures of **1** and **2** by 1- and 2-dimensional ^1^H, ^13^C, and ^15^N nuclear magnetic resonance (NMR) methods using heteronuclear multibond correlation (HMBC) and heteronuclear single quantum coherence (HSQC) spectroscopy. Analysis of ^1^H, ^1^H-^13^C HSQC and ^1^H-^13^C HMBC (2- and 3-bond C-H correlations) spectra of **1** assigned the two singlets at δ_H_ 7.83 and 7.54 to H-3 and H-6 of the trichlorophenyl ring; the aliphatic protons at δ_H_ 4.30 (t, *J*=6.4 Hz), 2.26 (m), and 4.16 (t, *J*=6.4 Hz) to the three CH_2_ groups at C-7 to C-9; and the six protons at δ_H_ 1.48 (s) to the two methyl groups at C-16 and C-16’ (Table S1, Fig. S6-13). Due to the lack of protons in the triazine ring and the four-bond separation between C-9 and the closest carbon, we were unable to unambiguously assign the chemical shifts for the triazine ring from ^1^H-^13^C data alone. We therefore recorded a ^1^H-^15^N HMBC (^2,3^*J*_HN_) experiment to assign the NH and NH_2_ chemical shifts and the regiochemistry of the ring. ^1^H-^15^N correlations from H-9, NH_2_-11’, NH-14 and Me-16/16’ to N-10 showed that the propoxy group must be bonded to N-10. ^1^H-^15^N correlations from Me-16/16’ and NH-14 to N-14, from NH-14 to N-13’, and from NH_2_-11’ to N-12 confirmed the composition of the triazine ring as shown in Fig. 2A. Nuclear Overhauser enhancements (NOEs) (Fig. S11) between H-9, Me-16/16’ and NH-14 provided additional proof of the N-O bond position. The structure of compound **1** was therefore as published for WR99210 (20, 21).

**Figure 2.**
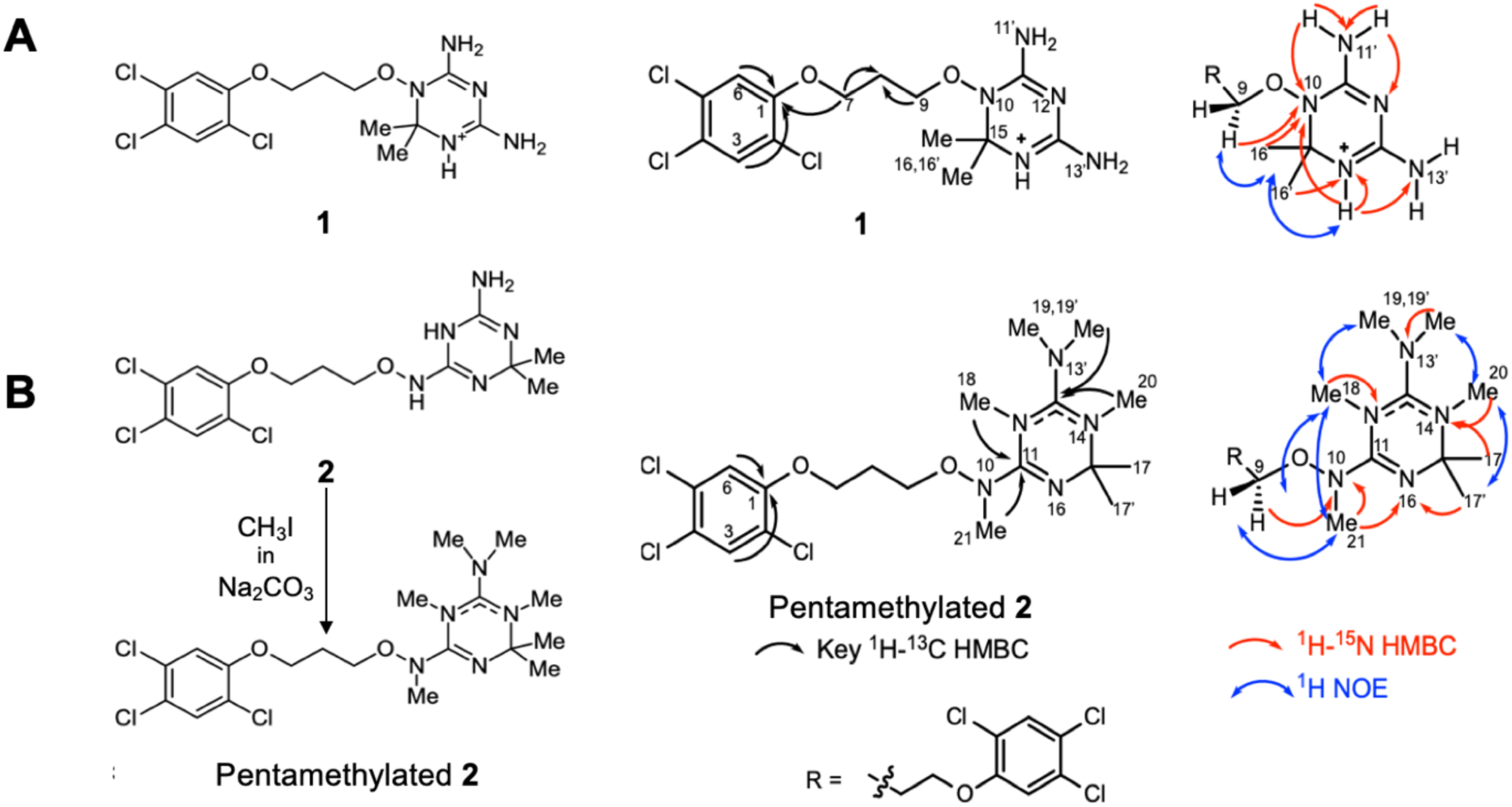
Structures of compounds 1 and 2 determined by nuclear magnetic resonance spectroscopy (NMR). (A) The structure of **1** was solved using ^1^H and ^15^N NMR without chemical derivatization. (B) Methylation of **2** introduced distinguishable hydrogens at the triazine ring, enabling definitive structural assignments. ^2,3^*J*_HC_ heteronuclear multiple bond correlation (HMBC) (black single side arrow); ^2,3^*J*_HN_ HMBC (red single side arrow), ^1^H-^15^N HMBC (red single side arrow), and ^1^H-^1^H NOE correlation (blue double-sided arrow).

NMR spectroscopy of compound **2** showed that the ^1^H and ^13^C NMR data corresponding to the (2,4,5-trichlorophenoxy) propoxy moiety, extending from C-1 through C-9, were in close agreement with **1**. However, the chemical shifts for the diamino-dimethyl triazine unit were absent, and the ^1^H spectrum showed broad peaks corresponding to two NH groups at δ_H_ 5.86 and 6.80 and one NH_2_ group at δ_H_ 5.61. Together, both the MS data and the NMR data suggested we had observed two, distinct singly-charged species by HRMS; namely a protonated and positively charged compound **2** or [**2**+H]^+^, and a positively charged compound **1**, or [**1**]^+^ already present in its protonated form (Fig. S2). The NMR data therefore demonstrate that **2** is an isomer of compound **1**. We recorded ^1^H-^15^N HSQC and HMBC spectra for **2** at multiple temperatures and in different solvents; however, unlike the case for compound **1**, we were unable to determine the complete structure of **2** by NMR alone (Table S2, Fig. S14-18).

To obtain additional NMR data we permethylated **2** using a stoichiometric excess of methyl iodide in sodium carbonate buffer (Fig. 2B). HRMS of the product indicated a molecular formula of C_19_H_29_Cl_3_N_5_O_2_, which corresponded to the addition of five methyl groups (Fig. S19). These were confirmed by signals from the ^1^H and ^13^C NMR spectra (Fig. S20-21, Table S3). The chemical shifts of all ^1^H, ^13^C and ^15^N atoms could be assigned from two-dimensional (2-D) ^1^H-^13^C and ^1^H-^15^N HSQC and HMBC data. The combination of ^1^H-^13^C and ^1^H-^15^N two- and three-bond correlations shown in black and red arrows, respectively, and the NOEs shown in blue arrows, allowed unambiguous assignment of the pentamethyl derivative of **2** as shown in Figure 2B. In particular, the HMBC correlations from H-9 and Me-21 to N-10, and from Me-21 and Me-17/17’ to N-16, together with NOEs between H-9 and Me-21, H-9 and Me-18, and Me-18 and methyl groups 19/19’ established the connectivity of the propoxy group to the triazine ring, thereby establishing the structure of compound **2** (Fig S22-25).

### Modeling of compounds 1 and 2 within the PfDHFR-TS binding pocket

Using the induced-fit docking mode of Glide, we examinined the predicted binding of compounds **1** and **2** into the active site of the PfDHFR-TS, as obtained from the known crystal structure of PfDHFR-TS complexed with WR99210 after removal of bound WR99210 (PDB ID: 1J3I, 2.33Å resolution). Glide iteratively combines ligand docking with protein remodeling around the binding site, and generates multiple poses of the receptor-ligand complex to compensate for to account for limitations in the scoring function (22, 23). Each of the poses employs unique structural modifications of the receptor for best fit, and these poses are ranked by GlideScore to predict the best structure of the docked complex.

The binding positions of WR99210 from the crystal structure and the poses of **1** in our model generated were evaluated by root mean square deviation (RMSD) calculations. Results confirmed that our model was in good agreement with the known binding geometry (Table S4). A RMSD of less than 1 Å is considered very good; the pose with the highest predicted stability gave a RMSD of 1.68 Å and the pose most similar to the crystalographic pose gave an RMSD of 0.61 Å (Table S4, Fig. 3A).

**Figure 3.**
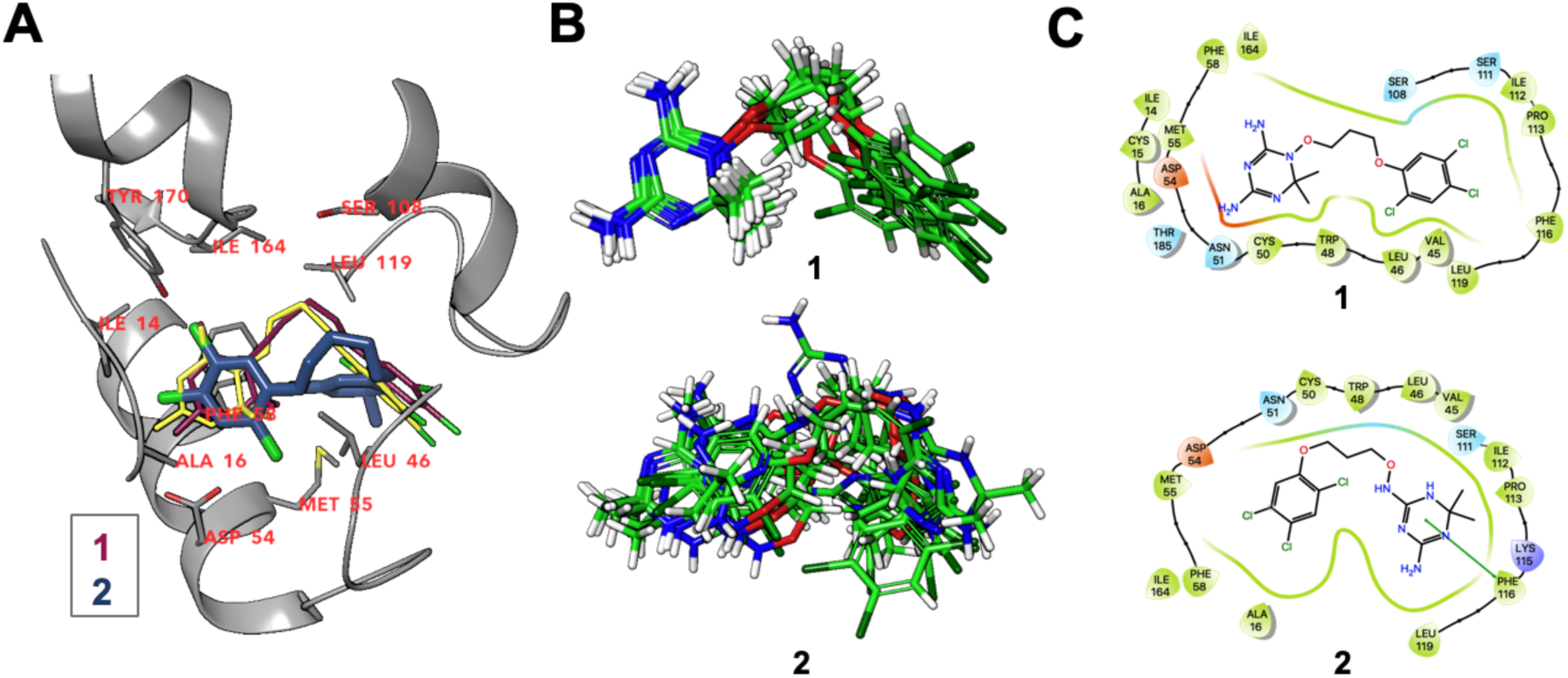
Crystal structure comparison, binding superposition, and Ligand Plots for compounds 1 and 2. (A) Overlay of the the crystal structure (yellow) and best-scoring predicted docked poses for each isomer shows **1** (purple) binds in a close agreement but **2** (blue) binds with opposite orientation, best visualized by comparison of the chlorine substituent position (green). (B) Superposition of the poses of **1** and **2** in the pocket. Results of docking **1** show overlapping poses as the favored thermodynamic solution, but results of docking with **2** return multiple disordered poses, without any favored arrangement. (C) Ligand Interaction Plot binding pocket models of 1 and 2 in the DHFR pocket showed pocket remodeling to accommodate **2.** A lack of favorable interactions between the amines on **2** and the pocket side chains for the isomer (represented as lines connecting the amino acid to the drug) is also observed. Interaction color code: hydrophobic, green; polar, blue; electrostatic negative side chain, red; electrostatic positive side chain, purple; π – π interactions, green line.

Docking scores loosely approximate binding energy, with more negative scores indicating greater stability. All of the poses generated for **1** had a better induced fit score than the best-scoring pose for **2**. Notably, all of the top docking poses of **1** had a binding mode similar to that of the known crystal structure (PDB ID: 1J3I), whereas the top scoring poses with **2** had numerous, dissimilar binding modes (Fig. 3B). The best-scoring docking pose with **1** had a GlideScore of −9.33 and an induced fit score of – 425.81, while the best-scoring pose of **2** produced a GlideScore of −8.07 and induced fit score of −423.44 (Fig. 3C). Furthermore, the best scoring binding pose of **2** was in an orientation opposite to that of **1** and the crystallographic pose, with the halogenated ring placed into the interior the enzyme rather than protruding from the pocket (Fig. 3A).

## DISCUSSION

Although not suitable for clinical use, WR99210 continues to be an important tool in molecular parasitology as a selection agent for genetically modified parasites. Recent observations of greatly different efficacies of WR99210 stocks from different sources have been unexplained. Here, we show that isomerization of WR99210 accounts for these efficacy differences, and we propose a rearrangement mechanism for isomerization. The potent antifolate activity of WR99210 that results from tight and specific binding to the *Plasmodium* DHFR-TS site is lost with the inactive isomer, which interacts only weakly at the DHFR active site in various, greatly different orientations.

HRMS analysis of WR99210 (compound **1**) and its isomer (compound **2**) showed that molecular formulae of these compounds were identical, without evidence in the stocks for degenerative products of reduced size or substantial impurities, incidental polymerization, or redox changes. HRMS also confirmed the experimental concentrations of **1** and **2** in DMSO-dissolved stocks used for drug response assays, thus demonstrating that the expected exposures of these compounds to parasitized erythroctyes were the same. Having eliminated these possible explanations for the large, supplier-dependent efficacy differences between the stocks, we performed TLC, RP-HPLC, and UV-vis spectroscopy studies on stocks of **1** and **2** for evidence of structural differences. All three methods identified differences between **1** and **2**. Compounds **1** and **2** migrated to different positions on TLC plates and were distinguished by their retention times in RP-HPLC. UV-vis spectroscopy showed clear differences in the absorption patterns of **1** and **2**, particularly in the 230 – 240 nanometer (nm) plateau region of the spectrum.

The structural solution of **2** and its elucidation by NMR as an isomer of **1** presented a challenge because of its paucity of distinguishable hydrogens on and in the immediate vicinity of the triazine ring. Methylation of the nitrogen atoms overcame this problem and provided the necessary signals for an NMR solution. The resulting structure of **2**, demonstrating that it is a positional isomer of **1**, is identical to that of a previous report, which demonstrated production of **2** from **1** in an ethanolic solution brought to pH >8 with sodium hydroxide, or with added triethylamine (21). Conversion of arylmethoxy-dimethyl-dihydrotriazines from their base form to isomeric dihydrotriazines had also been observed when the compounds were heated in ethanol, benzene, or partial aqueous suspension (24). Stocks of WR99210 may thus be inactivated by spontaneous rearrangement to isomer **2** under basic conditions.

Isomer **2** differs from WR99210 by a repositioning of the propoxyl substituent on the triazine ring. Figure 4 presents a proposed pathway of isomerization by a base-mediated ring opening, followed by ring closure at the gem dimethyl carbon with the NH_2_ group proximal to the propoxy linker. First, WR99210 forms from a substituted biguanide by a pathway involving *O*-bonded amine attack at the dimethyl-bearing methine carbon to form the triazine ring (8, 24, 25). Second, the amine groups of the diaminotriazine moiety in WR99210 are stabilized by the HCl salt form. When converted to, or purified as, the free base, tautomerization of the guanidine groups can occur, allowing for the prior imine nitrogen (now an NH_2_) to attack the dimethyl-bearing methine carbon to form the dihydrotriazine isomer **2**.

**Figure 4.**
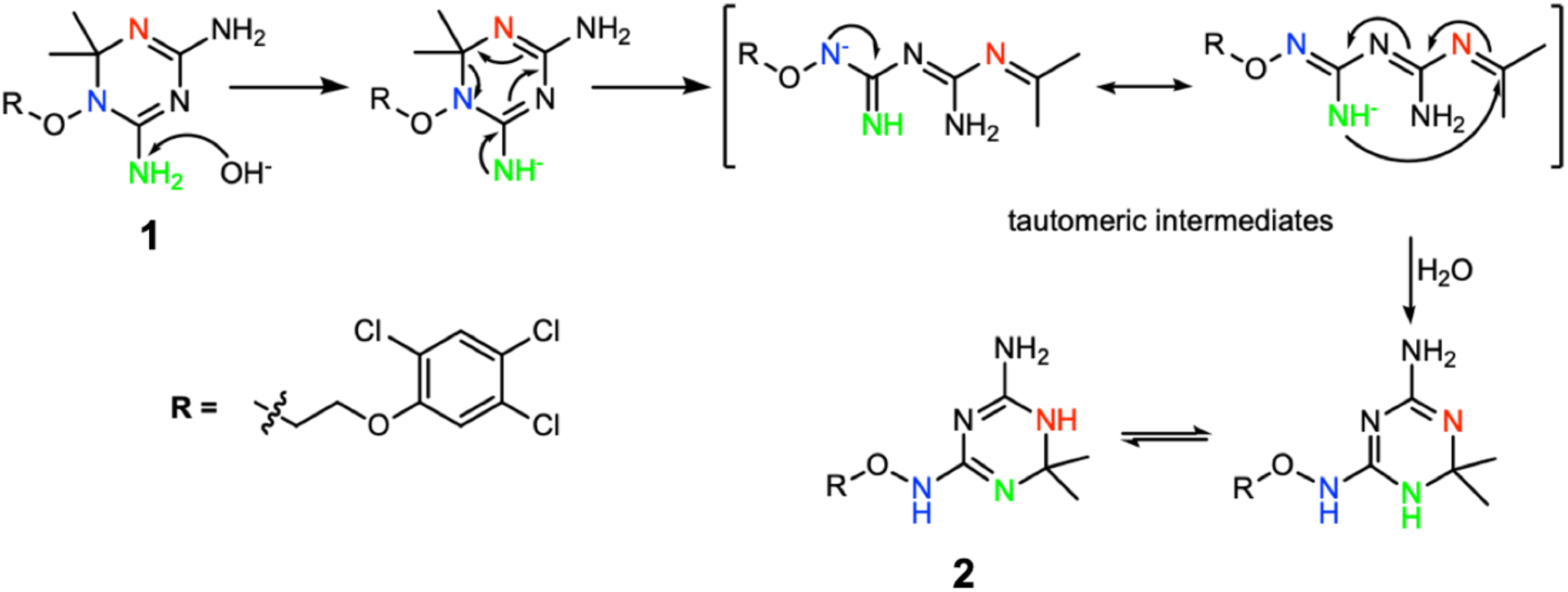
Proposed mechanism for the isomerization of compound 1 to 2. The mechanism of isomerization relies upon the conditions for production of the freebase rather than the hydrochloride salt, in agreement with the solved structures (Fig 2). The proposed mechanism starts with base-mediated ring opening, followed by ring closing via substitution at the gem dimethyl quaternary carbon by the NH_2_ group proximal to the propoxy linker. The isomerization which repositions the amine substituents and extends the propoxy linker between the halogenated ring and the triazine ring.

The results of our docking modeling indicate that isomer **2** binds to PfDHFR-TS with much lower affinity than WR99210, if at all. This dramatic difference of affinity further verifies the *Plasmodium* DHFR active site as WR99210’s target of action and accounts for the compound’s almost complete loss of drug efficacy after isomerization. Stocks and concentrates of WR99210 should therefore be checked for the presence of isomer **2**, particularly when they are not obtained and maintained as the hydrochloride salt or are exposed to basic conditions. Among the checks reported here are analysis by UV-vis spectroscopy, determination of the RP-HPLC elution time, and verification of the TLC band and its retention factor. We also note that infrared spectroscopy in the 6.0 – 10.0 µm region has been reported to distinguish diagnostic differences between unrearranged and rearranged triazines (24). These assessments alone or in combination may serve for accessible quality control of WR99210.

## MATERIALS AND METHODS

### Parasite Culture

*P. falciparum* Dd2 (Thailand) and NF54 (Netherlands) parasites were cultured in complete media (CM) consisting of iRPMI 1640 (KD Medical) buffered with 0.2% sodium bicarbonate (KD Medical). Media was additionally supplemented to 19.4 µg/ml gentamycin (KD Medical) and 1% AlbuMAX II (Gibco). Cultures were kept in a gassed incubator (5% O_2_, 5% CO_2_, 90% N_2_) at 37 °C. Initial morphology studies used synchronized wildtype NF54 strain cultures challenged with CM containing either 2 nM WR99210 (from JP or SA) or 1 μM WR99210 (SA). Sorbitol (5%, Sigma Aldrich) synchronization was performed 48 and 96 hrs prior to adding the drug media according to the published method (26). Media was changed and Giemsa-stained (Sigma Aldrich) thin smears were made daily.

### Solubilizing WR99210

<u>Dose response experiments</u> WR99210 stock solutions were resuspended with DMSO to 50 µM (JP) or 10 mM (SA) and stored at −20C. <u>Morphology experiments</u> Different lots of WR99210 either purchased from Sigma Aldrich (SA) or provided by Jacobus Pharmaceutical (JP), were dissolved in dimethylsulfoxide (DMSO, Sigma-Aldrich), and added to the media prior to sterile filtration. DMSO concentration was held the same in all test conditions (0.0002%), which had no observable effect on parasite development. HPLC, TLC, and MS experiments used a primary stock solution of 3.4 mg/ml WR99210 in tetrahydrofuran:methanol (2:1 vol/vol).

### Dose-response assays

*In vitro* drug response assessments were performed employing a standard 72-hour malaria SYBR Green I assay against the lab-adapted lines Dd2 and NF54 (27-29). Two-fold serial dilutions of JP and SA WR99210 (50 µL) were added across a 96-well plate, reserving two wells per row as drug-free controls. After reaching 4 – 10% parasitemia with >70% ring stage parasites, cultures were resuspended to 1% parasitemia and 1% hematocrit in CM and 150 µL was added to each well for drug phenotype response. EC_50_ values were determined using the variable sigmoidal function feature from Prism 7 on four independent replicates (GraphPad Software Inc).

### HPLC with UV-vis spectroscopy

A SB-C3, 3.5 micron, 300Å, 0.3 × 100 mm capillary column (Aligent) was used for chromatography at 6 ml/min. Typical sample amounts were 100 nanograms (ng). The column was equilibrated in 80% of 0.1% formic acid:20% acetonitrile and eluted over 15 minutes with a gradient to 100% acetonitrile. Detection was done at 290 nm with diode array spectral acquisition between 220 and 400 nm. Spectral analysis was performed with the Chemstation 2D software. For HPLC prior to Mass Spectrometry (further methods below) all chromatography conditions were maintained, except, a flow rate of 10 ml/min was used and approximately 25 ng of solubilized sample (primary stock solution) was injected.

### Mass Spectrometry

Acquisition and analysis was done with a Sciex 4000 QTrap in positive mode using either HPLC or direct infusion at 1 µg/ml of WR99210 in 85% acetonitrile:15% formic acid (0.1%) with a flow rate of 12 µl/min. Spray voltage was 4000, de-clustering potential was 25 volts and nebulizing nitrogen gas was 20 psi. MS2 fragmentation was done with a collision energy of 40 and MS3 with AFC values of 40 to 65.

### Nuclear Magnetic Resonance

NMR spectra were recorded on Bruker Avance 500 and 600 MHz spectrometers equipped with *z*-shielded gradient cryoprobes. Spectra were recorded at multiple temperatures, and 1- and 2-dimensional homonuclear and heteronuclear ^1^H, ^13^C and ^15^N spectra were recorded. See Supporting Information for additional details.

### Computational docking studies

The three-dimensional structure of PfDHFR-TS (PDB ID 1J3I) was downloaded from the Protein Data Bank. The protein structure was readied for docking via the Protein Preparation procedure, and used Induced Fit Docking protocol 2015-2, Glide version 6.4, Prime version 3.7, Schrödinger, LLC, New York, NY, 2015, release 2018-4.

To achieve unbiased docking, the conformation of WR99210 from the PDB structure was not used. Instead, the 3D conformer of WR99210 was downloaded from PubChem using CID 121750. The SA isomer was drawn in the PubChem sketcher and converted to 3D using Open Babel 2.3.1 (30). Both ligands were readied for docking using the LigPrep procedure in Maestro (31).

Induced fit docking (IFD) was performed using the Maestro suite. IFD used Glide for ligand docking with a softened potential to increase the possible initial protein conformations by decreasing the van der Waals repulsion term, permitting closer packing, and a final re-docking after protein optimization. Prime was used to predict side-chain position within each protein-ligand complex and to find stable coformations around the docked ligand. To evaluate the quality of the docked poses, IFD provided an energy score for each docked pose that combines both Glide and Prime. Root mean square deviation (RMSD) calculations were performed between the crystallographic structure and the poses resulting from IFD using the DockRMSD program (https://zhanglab.ccmb.med.umich.edu/DockRMSD/). 2D ligand interaction diagrams of the active site were created within Maestro and virtual reality visualization was done via UCSF ChimeraX (https://www.rbvi.ucsf.edu/chimerax/) and the HTC Vive Pro.

## ACKNOWLEDGEMENTS

We thank Joseph Brzostowski for his assistance with microscopy, Anna K. Crater for advice and discussions, Anna Liu for her help with EC_50_ plates, and Jianbing Mu for providing *P. falciparum* NF54 and generous advice during the project.

Virtual reality graphics and analyses performed with UCSF ChimeraX, developed by the Resource for Biocomputing, Visualization, and Informatics at the University of California, San Francisco, with support from National Institutes of Health R01-GM129325 and the Office of Cyber Infrastructure and Computational Biology, National Institute of Allergy and Infectious Diseases.

## Author contributions

KDL, TPR, CAB, SDG, and TEW designed the experiments; KDL did the drug response experiments and analyzed the data; TPR did the morphology assessments; GAN did the TLC and HPLC experiments; GAN, GH, and SDG did the MS experiments; SDG and CAB designed the NMR experiments and SDG, CAB, and RDO performed the experiments and analyzed the data; PC designed and performed the docking experiments; TPR, SDG, CAB, TEW, and KDL wrote the manuscript; all authors edited and approved the manuscript.

This research was supported by the Intramural Research Program of the NIH, National Institute of Allergy and Infectious Diseases (NIAID) and the National Institute of Diabetes and Digestive and Kidney Diseases (NIDDK).

GH is an employee of Jacobus Pharmaceutical Company, Inc.

