## Supplemental files for "Isomerization of antimalarial drug WR99210 explains its inactivity in a commercial stock"

### Supplemental Information

**Supplemental Methods.** Microscopy, TLC, Chemistry and structure determination.
**Table S1.** $^{1}$H (500 MHz) and $^{13}$C (125 MHz) NMR data of compound **1** in DMSO-$d_6$
**Table S2.** $^{1}$H (500 MHz) and $^{13}$C (125 MHz) NMR data of compound **2** in DMSO-$d_6$
**Table S3.** $^{1}$H (500 MHz) and $^{13}$C (125 MHz) NMR data of compound **3** in DMSO-$d_6$
**Table S4.** Computational Modeling Scores of **1** and **2** bound to PfDHFR-TS
**Figure S1.** HRMS spectrum of **1**.
**Figure S2.** HRMS spectrum of **2**.
**Figure S3.** TLC of **1** and **2**
**Figure S4.** HPLC of **1**
**Figure S5.** HPLC of **2**
**Figure S6.** $^{1}$H NMR spectrum of **1** in DMSO-$d_6$ at 328K.
**Figure S7.** $^{13}$C NMR spectrum of **1** in DMSO-$d_6$ at 298K.
**Figure S8.** $^{1}$H-$^{13}$C HSQC spectrum of **1** in DMSO-$d_6$ at 328K.
**Figure S9.** $^{1}$H-$^{13}$C HMBC spectrum of **1** in DMSO-$d_6$ at 328K.
**Figure S10.** $^{1}$H-$^{15}$N HSQC coupled spectrum of **1** in DMSO-$d_6$ at 298K.
**Figure S11.** $^{1}$H-$^{15}$N LR-HSQMBC of **1** in DMSO-$d_6$ at 298K.
**Figure S12.** $^{1}$H-$^{15}$N LR-HSQMBC spectrum of **1** in DMSO-$d_6$ at 328K.
**Figure S13.** 1-D $^{1}$H NOE spectrum of **1** in DMSO-$d_6$ at 328K.
**Figure S14.** $^{1}$H NMR spectrum of **2** in DMSO-$d_6$ at 298K.
**Figure S15.** $^{13}$C NMR spectrum of **2** in DMSO-$d_6$ at 298K.
**Figure S16.** $^{1}$H-$^{13}$C HSQC coupled spectrum of **2** in DMSO-$d_6$ at 298K.
**Figure S17.** $^{1}$H-$^{13}$C HMBC coupled spectrum of **2** in DMSO-$d_6$ at 298K.
**Figure S18.** $^{1}$H-$^{15}$N LR-HSQMBC coupled spectrum of **2** in DMSO-$d_6$ at 298K.
**Figure S19** HRMS spectrum of **3**.
**Figure S20.** $^{1}$H NMR spectrum of **3** in DMSO-$d_6$ at 297.9K.
**Figure S21.** $^{13}$C NMR spectrum of **3** in DMSO-$d_6$ at 298K.
**Figure S22.** $^{1}$H-$^{13}$C HSQC coupled spectrum of **3** in DMSO-$d_6$ at 298K.
**Figure S23.** $^{1}$H-$^{13}$C HMBC coupled spectrum of **3** in DMSO-$d_6$ at 298K.
**Figure S24.** $^{1}$H-$^{15}$N HMBC coupled spectrum of **3** in DMSO-$d_6$ at 298K.
**Figure S25.** 1-D $^{1}$H NOE spectrum of **3** in DMSO-$d_6$ at 298K.

**Supplemental Experimental Methods**

**Microscopy.** Giemsa-stained thin smears of cultures were fixed for 1 minute in 100% methanol (MG Scientific) and stained in 10% Giemsa stain for 15 minutes after which they were subsequently rinsed with distilled water and allowed to air dry. Images were taken on an upright, Zeiss Axio Imager A1 microscope using Zeiss AxioVision software and employing a Plan-FLUAR 100X, 1.45 oil-immersion objective.

**TLC.** 5 × 10 cm glass backed silica gel plates (Sigma Aldrich) were developed with chloroform:diethyl ether:methanol (5:3:2 vol/vol) and visualized by UV light. Sample spotting volumes (tetrahydrofuran:methanol, 2:1) were 1 μl and contained approximately 1 μg of each dissolved compound.

**Chemistry and structure determination.** Compound **1** was obtained from Jacobus Pharmaceutical Company, Inc. and compound **2** from Sigma-Aldrich. All other reagents and HPLC-grade solvents used in these studies were obtained from Sigma-Aldrich. HPLC purification was performed on a semi-preparative HPLC (Agilent 1160) using a Waters Symmetry Prep™ C18 column (7.8 × 300 mm, 7 μm). NMR spectra were recorded on a Bruker Avance500 spectrometer using a triple resonance cryoprobe and *z* gradients. Spectra were recorded at multiple temperatures. Data recorded at 298 K and 328 K are included here.

Compound **3** (pentamethyl derivative of compound **2**): 4 mg (10 μmol) of **2** obtained from Sigma Aldrich was dissolved in DMSO (100 μL) in the presence of $Na_2CO_3$ (2.6 mg) and MeI (2.2 μL, 35 μM) and stirred at rt for 16 h. The resulting mixture was diluted with water (1 mL) and purified by semipreparative HPLC (flow rate of 3 mL/min, eluting with a linear gradient of 0.5–20%, $CH_3CN$ in $H_2O$ with 0.1% TFA in 30 min) to give compound **3** (1.5 mg, $t_R$ 10.6 min). Structure determination of **3** was accomplished by interpretation of HRMS, and 1D and 2D NMR experiments (see below) and proved the structure of **2**.

**Table S1.** $^{1}H$ (500 MHz) and $^{13}C$ (125 MHz) NMR data of compound **1** in DMSO-$d_6$

| Atom | $\delta_C$, type[a] | $\delta_N$, type[b] | $\delta_H$, mult (*J* in Hz)[c] |
|---|---|---|---|
| 1 | 153.2, C | | |
| 2 | 130.6, C | | |
| 3 | 130.6, CH | | 7.83, s |
| 4 | 122.8, C | | |
| 5 | 121.1, C | | |
| 6 | 115.5, CH | | 7.53, s |
| 7 | 73.9, $CH_2$ | | 4.30, t (6.4) |
| 8 | 26.8, $CH_2$ | | 2.26, m |
| 9 | 65.9, $CH_2$ | | 4.16, t (6.4) |
| 10 | | 175, N | |
| 11 | 161.2, C | | |
| 11’ | | 94, $NH_2$ | 8.05, s 8.61, s |
| | | | 8.61, s |
| 12 | | 151, N | |
| 13 | 156.3, C | | |
| 13’ | | 87, $NH_2$ | 7.54, m |
| 14 | | 106, NH | 9.41, s |
| 15 | 72.2, C | | |
| 16[d] | 25.0, $CH_3$ | | 1.48, s |
| 16’[d] | 22.3, $CH_3$ | | 1.48, s |

[a]Recorded at 125 MHz; referenced to residual DMSO-$d_6$, at δ 39.5 ppm.
[b]Recorded at 50 MHz; referenced via the $^{15}N$ gyromagnetic ratio
[c]Recorded at 500 MHz; referenced to residual DMSO-$d_6$, at δ 2.50 ppm
[d]Can be interchanged.

**Table S2.** $^{1}H$ (500 MHz) and $^{13}C$ (125 MHz) NMR data of compound **2** in DMSO-$d_6$

| Atom | $\delta_C$, type[a] | $\delta_N$, type[b] | $\delta_H$, mult (*J* in Hz)[c] |
|---|---|---|---|
| 1 | 154.0, C | | |
| 2 | 131.08, C | | |
| 3 | 131.05, CH | | 7.86, s |
| 4 | 122.9, C | | |
| 5 | 121.5, C | | |
| 6 | 115.8, CH | | 7.51, s |
| 7 | 67.5, $CH_2$ | | 4.24, t (6.4) |
| 8 | 28.8, $CH_2$ | | 2.07, m |
| 9 | 68.2, $CH_2$ | | 3.89, t (6.4) |
| 10 | | 95, NH | 6.80[d] |
| 11 | 154.6, C | | |
| 12 | | na,[e] N | |
| 13 | 155.0, C | | |
| 13’ | | 71, $NH_2$ | 5.61, br s |
| 14 | | 83, NH | 5.86[d] |
| 15 | 63.9, C | | |
| 16 | | na,[e] N | |
| 17,17’ | 29.7, $CH_3$ | | 1.31, s |

[a]Recorded at 125 MHz; referenced to residual DMSO-$d_6$, at δ 39.5 ppm
[b]Recorded at 50 MHz; referenced via the $^{15}N$ gyromagnetic ratio
[c]Recorded at 500 MHz; referenced to residual DMSO-$d_6$, at δ 2.50 ppm
[d]Signals can be interchanged
[e]na, not assigned

**Table S3.** $^{1}H$ (500 MHz) and $^{13}C$ (125 MHz) NMR data of compound **3** in DMSO-$d_6$

| Atom | $\delta_C$, type[a] | $\delta_N$, type[b] | $\delta_H$, mult ($J$ in Hz)[c] |
|---|---|---|---|
| 1 | 152.8, C | | |
| 2 | 131.0, C | | |
| 3 | 131.1, CH | | 7.90, s |
| 4 | 123.3, C | | |
| 5 | 121.6, C | | |
| 6 | 115.9, CH | | 7.54, s |
| 7 | 66.4, $CH_2$ | | 4.25, t (6.1) |
| 8 | 27.9, $CH_2$ | | 2.11, m |
| 9 | 68.1, $CH_2$ | | 4.12, t (6.1) |
| 10 | | 157, N | 3.00, s |
| 11 | 153.7, C | | |
| 12 | | 112, N | 3.36, s |
| 13 | 158.6, C | | |
| 13’ | | 69, N | 2.96, s |
| 14 | | 111, N | 3.11, s |
| 15 | 73.9, C | | |
| 16 | | 241, N | |
| 17/17’ | 27.8, $CH_3$ | | 1.48, s |
| 18 | 37.9, $CH_3$ | | 3.36, s |
| 19/19’ | 40.7, $CH_3$ | | 2.96, s |
| 20 | 35.0, $CH_3$ | | 3.11, s |
| 21 | 37.3, $CH_3$ | | 3.00, s |

[a]Recorded at 125 MHz; referenced to residual DMSO-$d_6$, at δ 39.5 ppm
[b]Recorded at 50 MHz; referenced via the $^{15}N$ gyromagnetic ratio
[c]Recorded at 500 MHz; referenced to residual DMSO-$d_6$, at δ 2.50 ppm

**Table S4.** Using the induced-fit docking mode of Glide, we examined the predicted binding of compounds 1 and 2 into the active site of the PfDHFR-TS, as obtained from the known crystal structure solution of PfDHFR-TS complexed with WR99210 after removal of bound WR99210 (PDB ID: 1J3I, 2.33Å resolution).

**Table S4.** Computational Modeling Scores of **1** and **2** bound to PfDHFR-TS

| Compound | Docking Score | Induced Fit Docking Score | Best Pose RMSD | Lowest RMSD* |
|---|---|---|---|---|
| JP WR99210 **(1)** | -9.33 | -425.81 | 1.68 Å | 0.61 Å |
| SA WR99210 **(2)** | -8.07 | -423.44 | N/A | N/A |

The best scoring poses are reported in all cases (except in * which is the lowest RMSD value generated), study conducted using PDB ID: 1J3I, RMSD = Root Mean Square Deviation - uses the drug position crystal structure as the reference and cannot be calculated for SA due to a lack of equivalent atoms between the isomers

**Figure S1.** HRMS spectrum of **1**.

### Elemental Composition Report



#### Single Mass Analysis

Tolerance = 5.0 mDa / DBE: min = -1.5, max = 100.0
Element prediction: Off
Number of isotope peaks used for i-FIT = 3

Monoisotopic Mass, Even Electron Ions
116 formula(e) evaluated with 2 results within limits (up to 50 closest results for each mass)
Elements Used:
C: 0-100 H: 0-250 N: 3-5 O: 0-60 35Cl: 3-3
SG-19DEC19-4-S1-JP 102 (1.742) AM2 (Ar,25000.0,0.00,0.00); ABS
TOF MS ES+

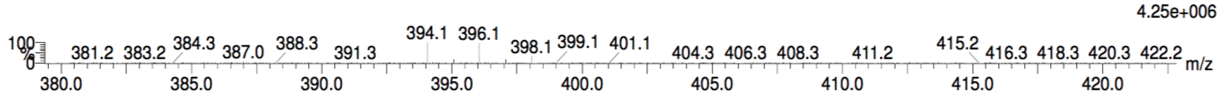


| | | | | | | | | |
|---|---|---|---|---|---|---|---|---|
| Minimum: | | | | -1.5 | | | | |
| Maximum: | | 5.0 | 5.0 | 100.0 | | | | |
| Mass | Calc. Mass | mDa | PPM | DBE | i-FIT | Norm | Conf(%) | Formula |
| 394.0606 | 394.0604 | 0.2 | 0.5 | 6.5 | 539.7 | 0.492 | 61.17 | C14 H19 N5 O2 35Cl3 |
| | 394.0645 | -3.9 | -9.9 | 10.5 | 540.2 | 0.946 | 38.83 | C19 H19 N3 35Cl3 |

**Figure S2.** HRMS spectrum of **2**.

### Elemental Composition Report



#### Single Mass Analysis

Tolerance = 5.0 mDa / DBE: min = -1.5, max = 100.0
Element prediction: Off
Number of isotope peaks used for i-FIT = 3

Monoisotopic Mass, Even Electron Ions
116 formula(e) evaluated with 2 results within limits (up to 50 closest results for each mass)
Elements Used:
C: 0-100 H: 0-250 N: 3-5 O: 0-60 35Cl: 3-3
SG-19DEC19-4-S1-SA 136 (2.317) AM2 (Ar,25000.0,0.00,0.00); ABS
TOF MS ES+

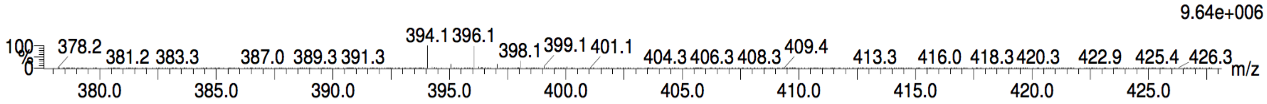


| | | | | | | | | |
|---|---|---|---|---|---|---|---|---|
| Minimum: | | | | -1.5 | | | | |
| Maximum: | | 5.0 | 5.0 | 100.0 | | | | |
| Mass | Calc. Mass | mDa | PPM | DBE | i-FIT | Norm | Conf(%) | Formula |
| 394.0603 | 394.0604 | -0.1 | -0.3 | 6.5 | 578.6 | 0.487 | 61.45 | C14 H19 N5 O2 35Cl3 |
| | 394.0645 | -4.2 | -10.7 | 10.5 | 579.0 | 0.953 | 38.55 | C19 H19 N3 35Cl3 |

**Figure S3.** TLC of **1** and **2.** A solvent system of chloroform:diethyl ether:methanol, (5:3:2 vol/vol) was used to separate the compounds which were then visualized under UV light. Lanes, from left to right: **2, 2** separate lot, **1,** co-spot of **1** and **2** first lot, co-spot of **1** and **2** second lot.

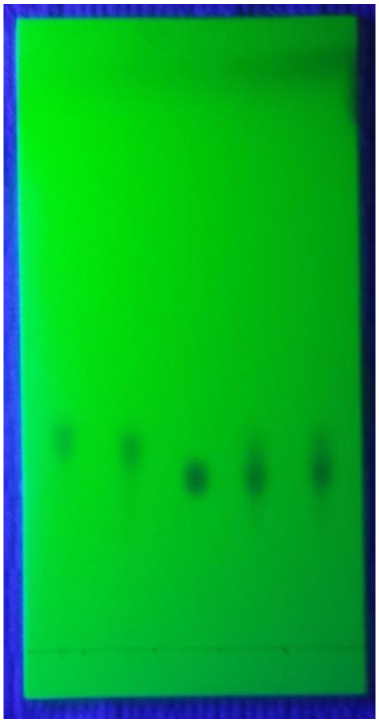

**Figure S4.** HPLC of **1.** With the monitoring wavelength set at 290 nM, **1** gave a retention time of 13.7 min. See supplemental methods for experimental detail.

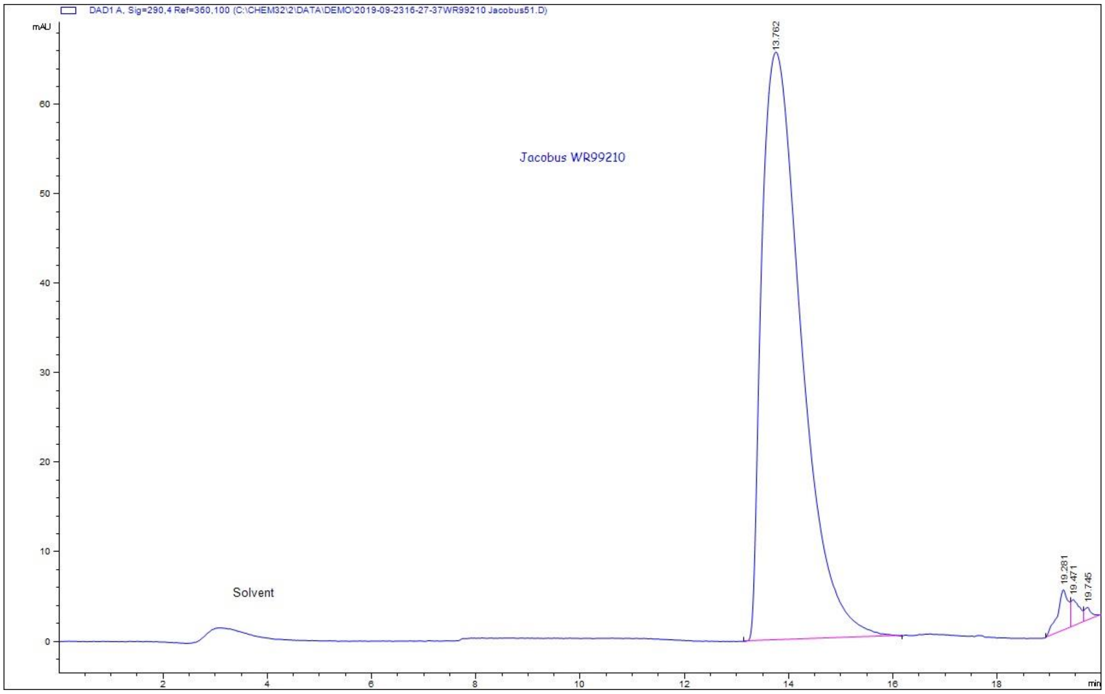

**Figure S5.** HPLC of **2.** With the monitoring wavelength set at 290 nM, **2** gave a retention time of 13.1 min. See supplemental methods for experimental detail.

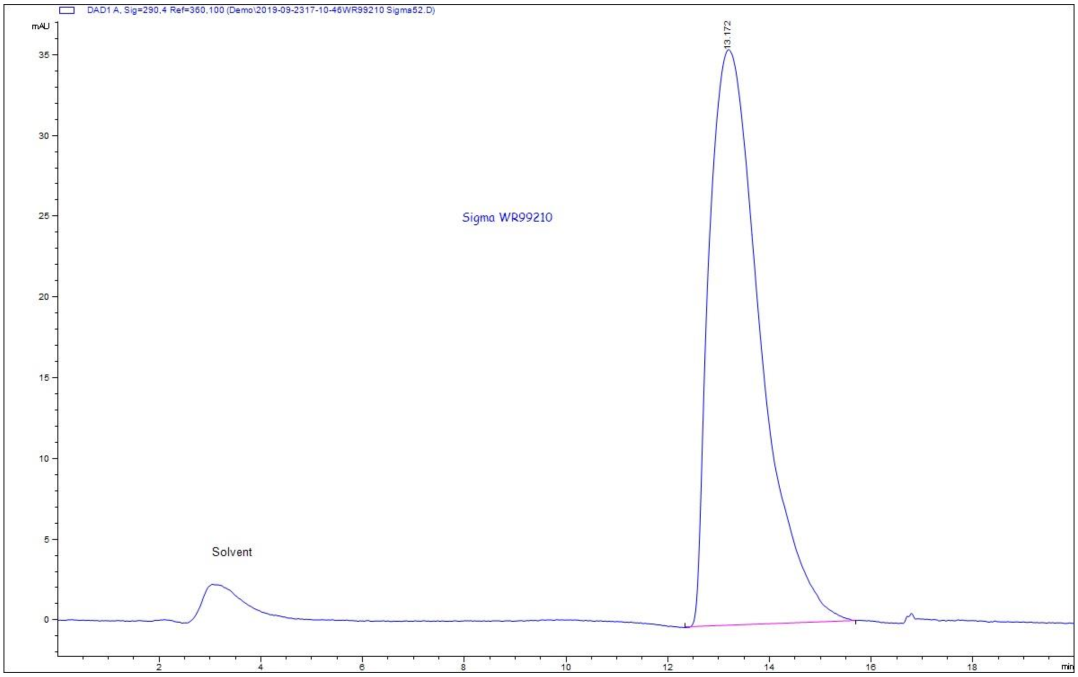

**Figure S6.** $^{1}$H NMR spectrum of 1 in DMSO-$d_6$ at 328K.

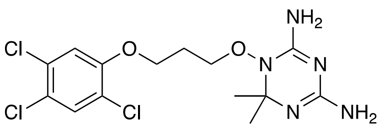

**1**

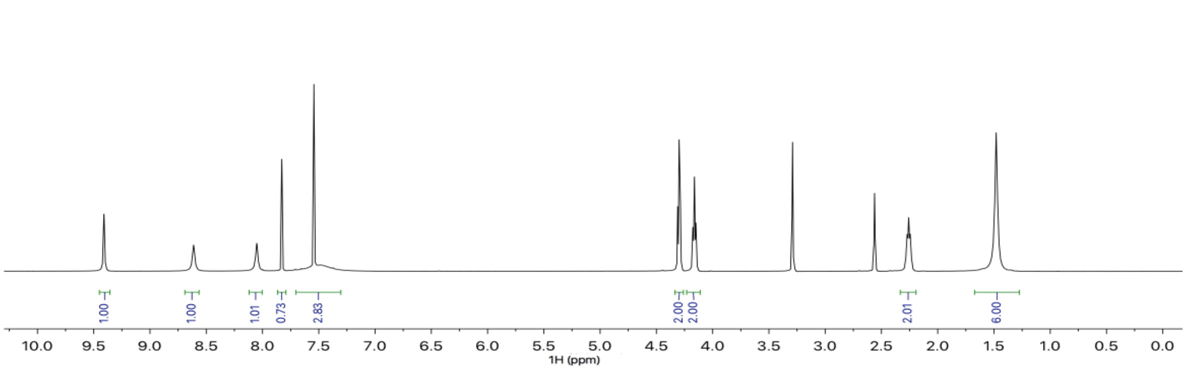

**Figure S7.** $^{13}$C NMR spectrum of **1** in DMSO-$d_6$ at 298K.

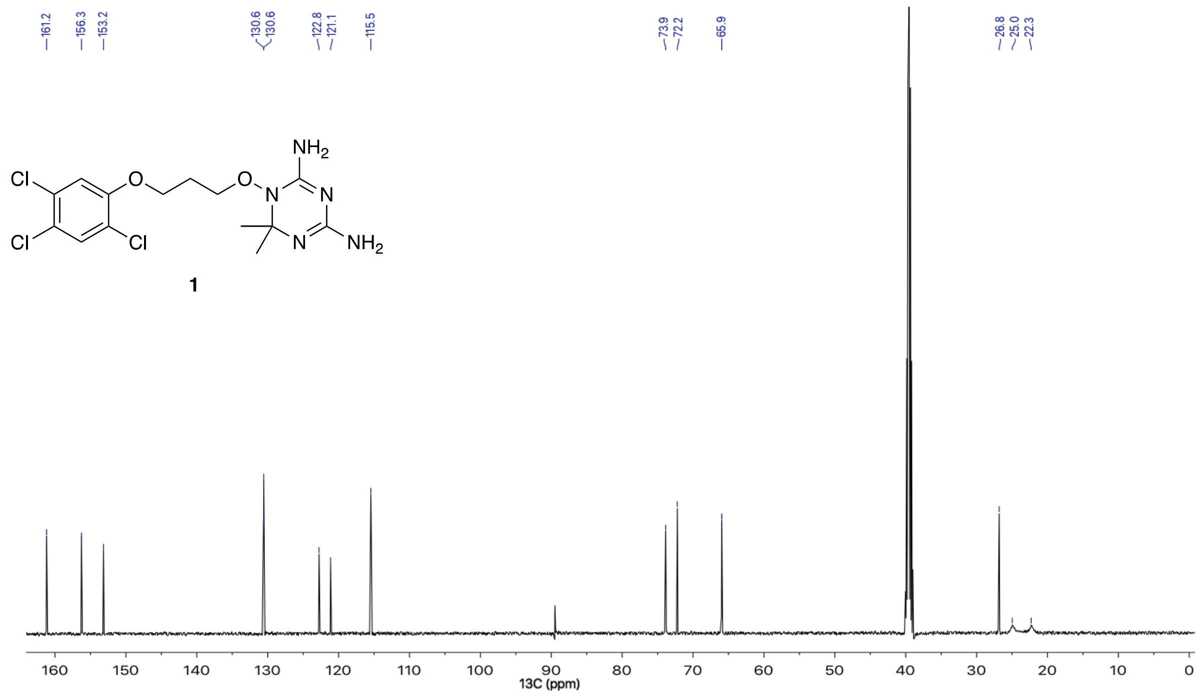

**Figure S8.** $^1H$-$^{13}C$ HSQC spectrum of **1** in DMSO-$d_6$ at 328K.

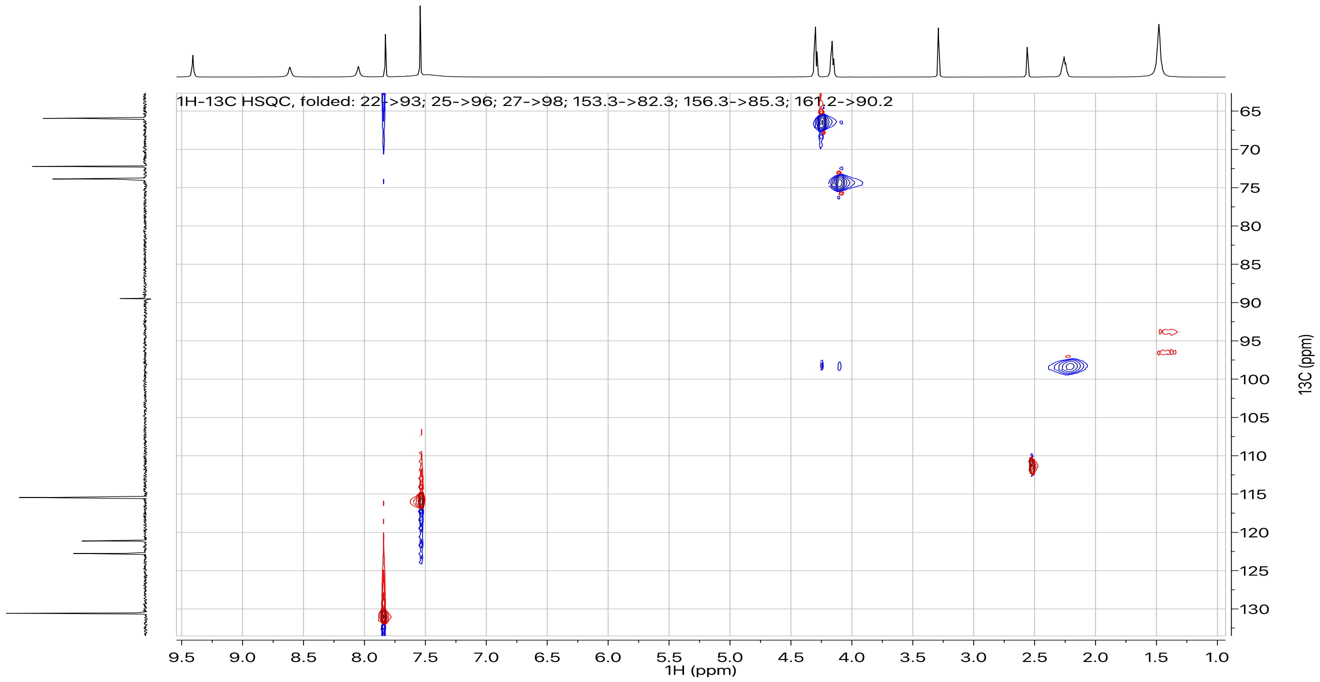

**Figure S9.** $^{1}$H-$^{13}$C HMBC spectrum of **1** in DMSO-$d_6$ at 328K.
Recorded with a $^{13}$C sweep width of 71 ppm and the carrier (O2P) set at 98 ppm; folded peaks are labeled on the spectral heading.

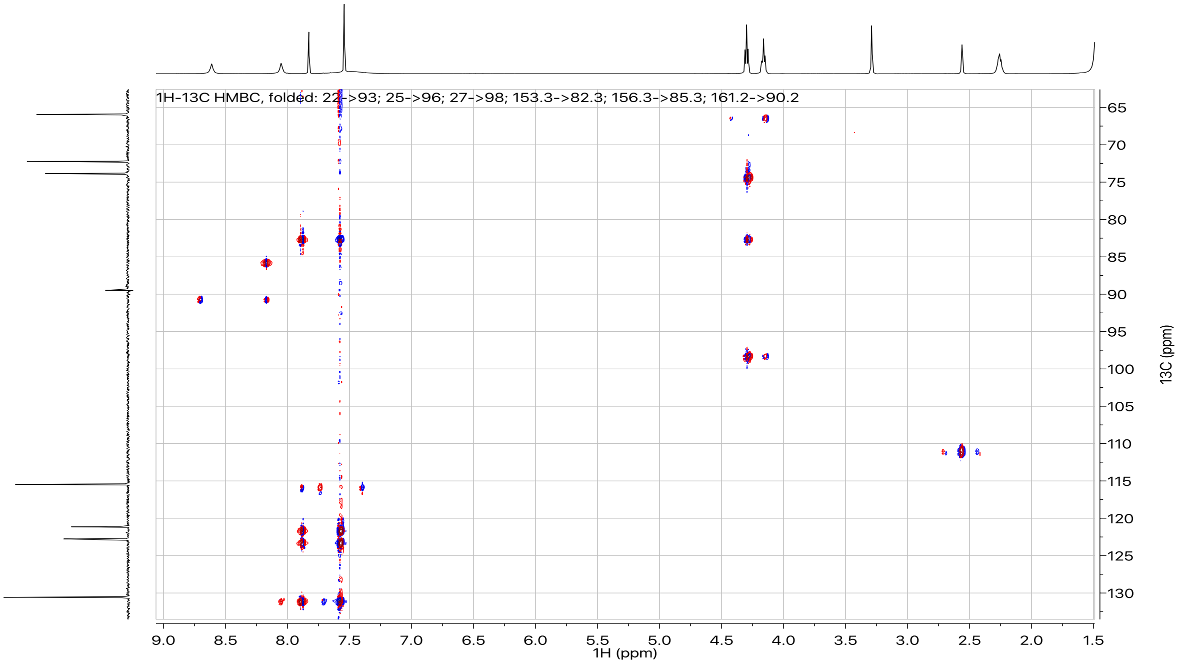

**Figure S10.** $^{1}H$-$^{15}N$ HSQC coupled spectrum of **1** in DMSO-$d_6$ at 298K.

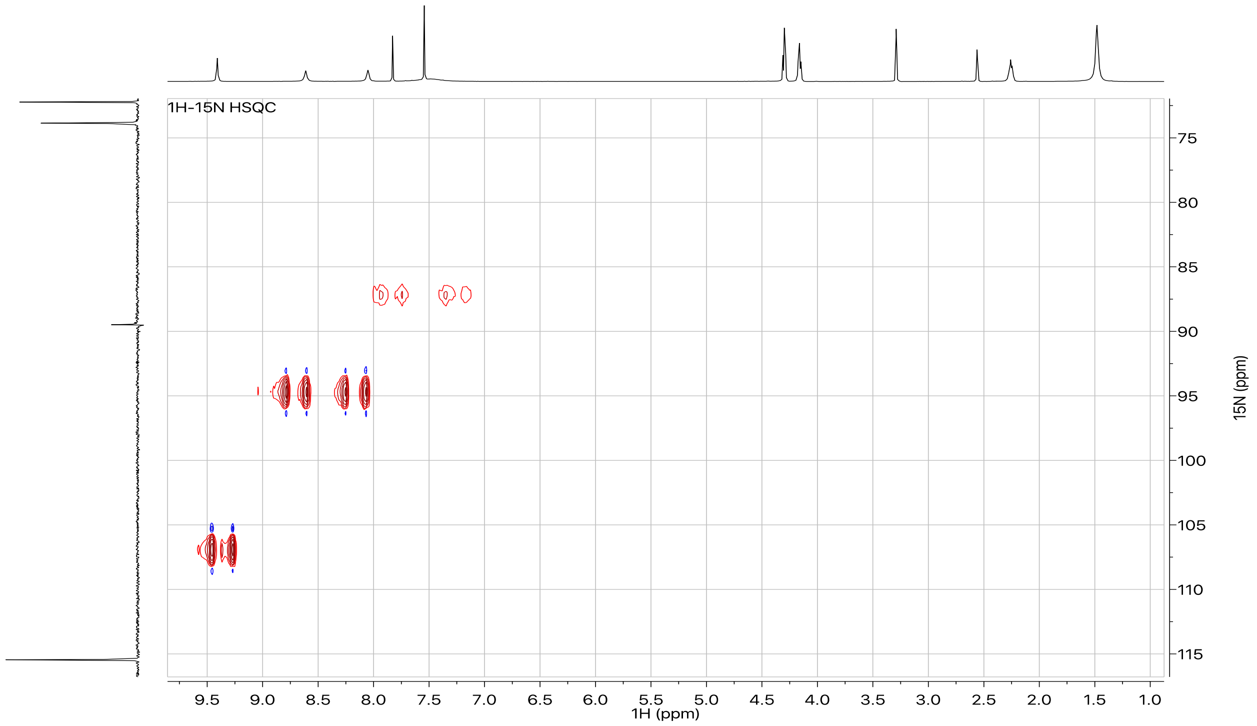

**Figure S11.** $^{1}H$-$^{15}N$ LR-HSQMBC spectrum of **1** in DMSO-$d_6$ at 298K.

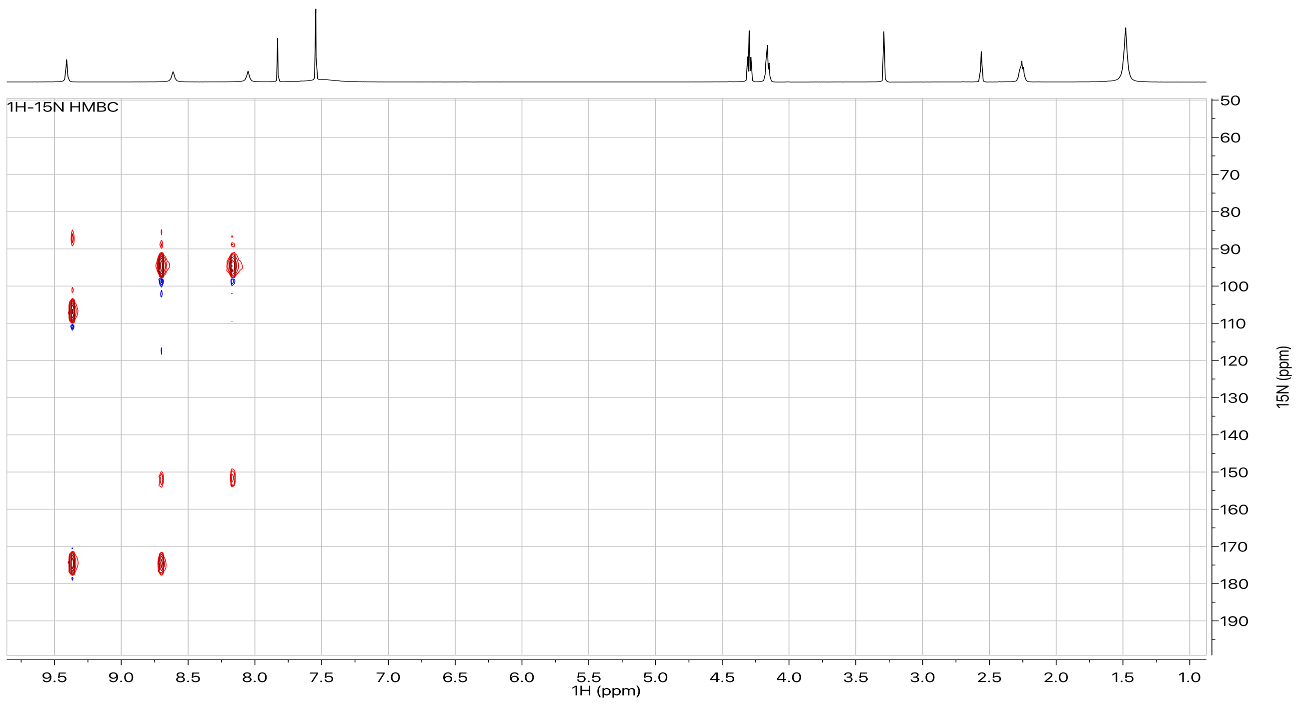

**Figure S12.** $^{1}H$-$^{15}N$ LR-HSQMBC spectrum of **1** in DMSO-$d_6$ at 328K.

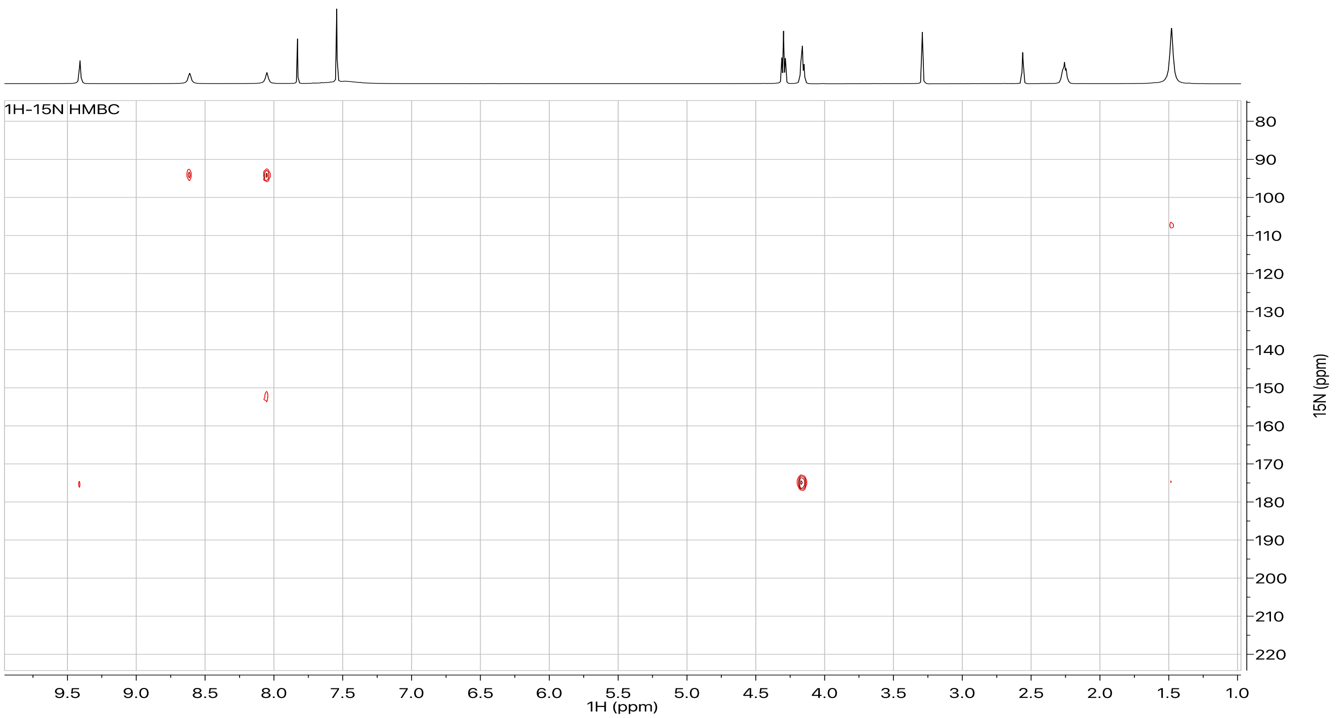

**Figure S13.** 1-D $^{1}H$ NOE spectrum of **1** in DMSO-$d_6$ at 328K. $^{1}H$'s that were irradiated in each spectrum are labeled on the figure.

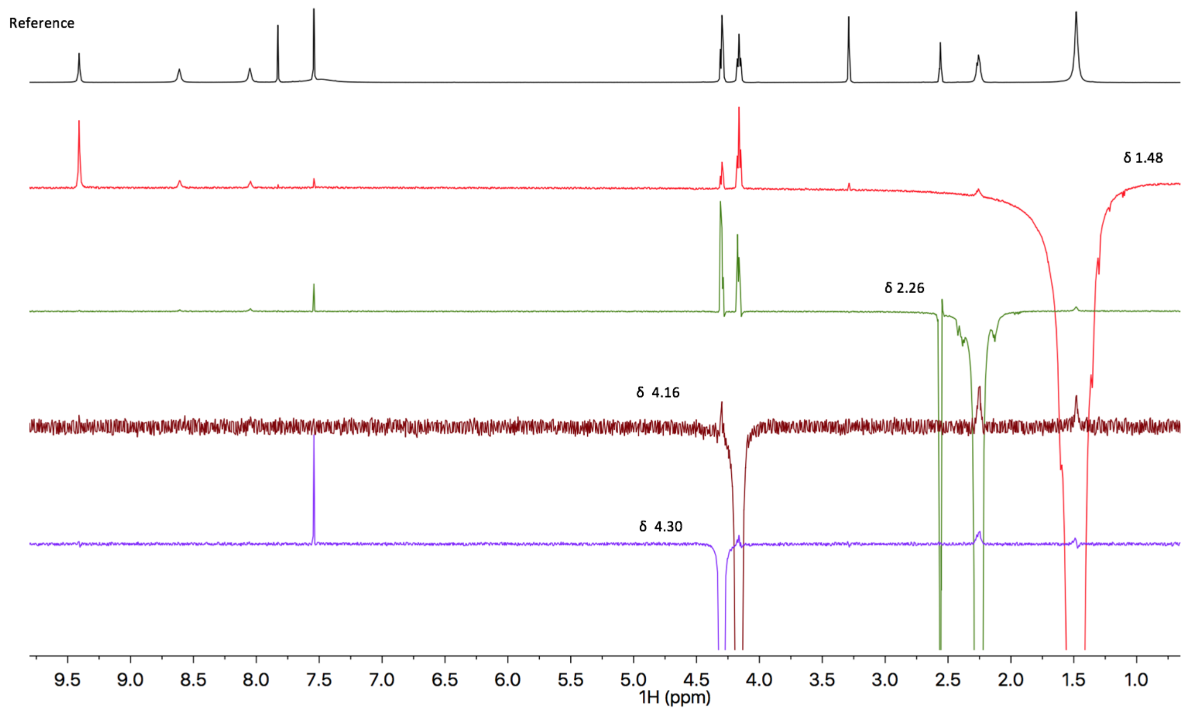

**Figure S14.** $^{1}$H NMR spectrum of **2** in DMSO-$d_6$ at 298K.

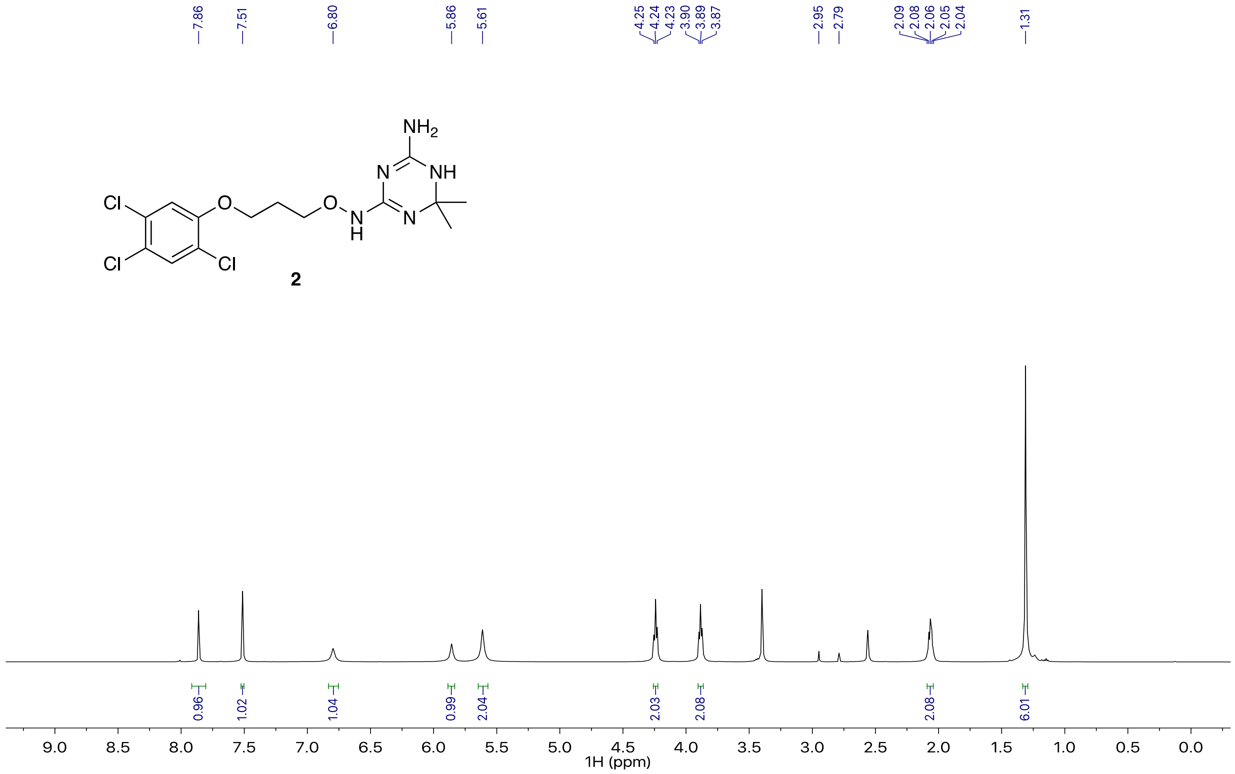

**Figure S15.** $^{13}$C NMR spectrum of **2** in DMSO-$d_6$ at 298K.

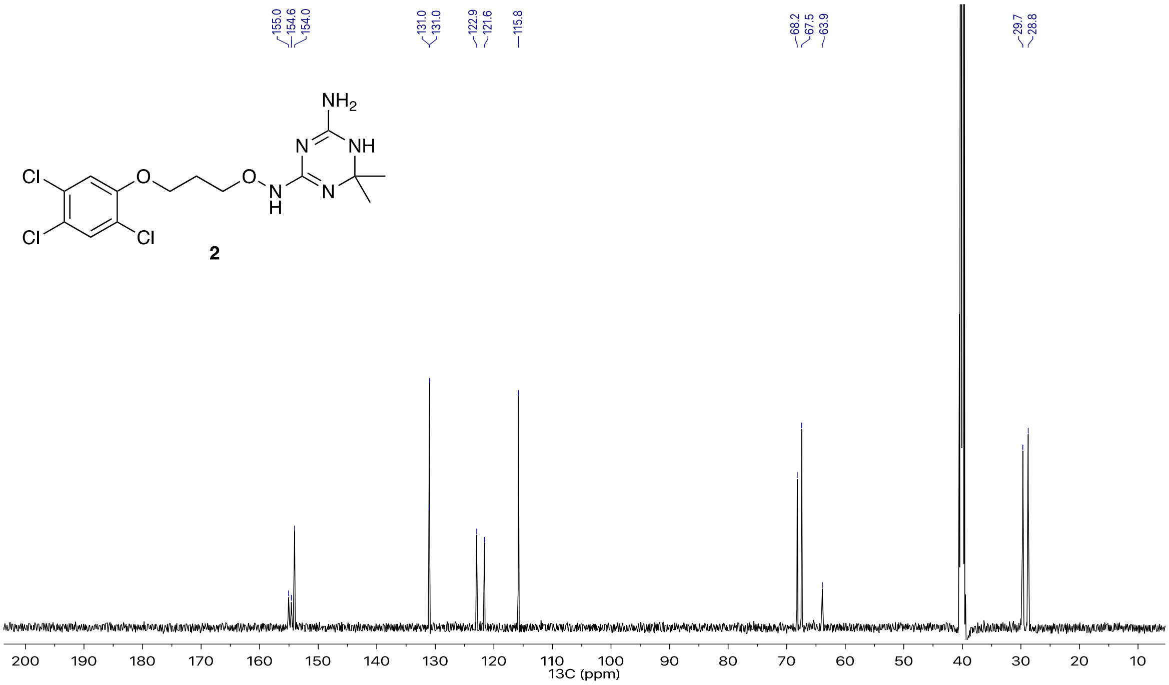

**Figure S16.** $^{1}$H-$^{13}$C HSQC coupled spectrum of **2** in DMSO-$d_6$ at 298K.

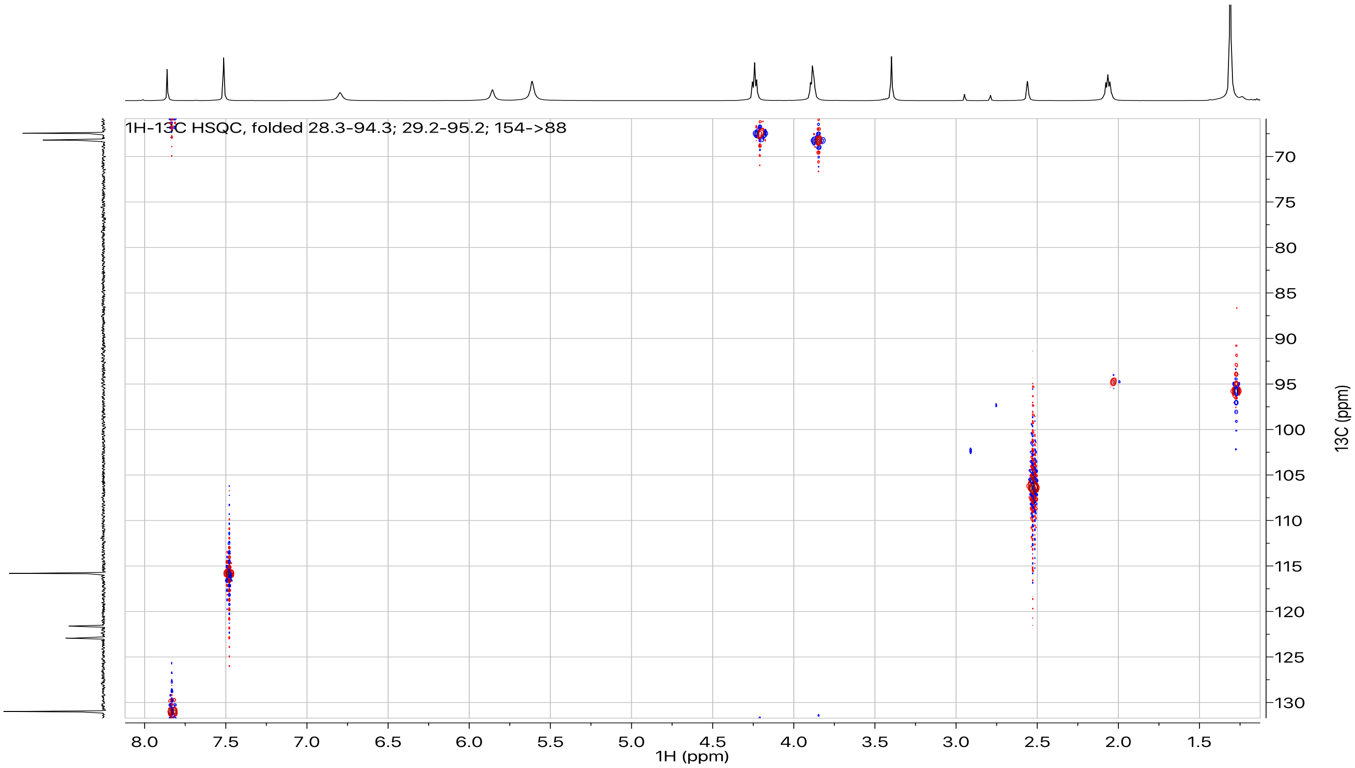

**Figure S17.** $^1$H-$^{13}$C HMBC coupled spectrum of **2** in DMSO-$d_6$ at 298K.
Recorded with a $^{13}$C sweep width of 72 ppm and the carrier set at 98 ppm; folded peaks are labeled on the spectral heading.

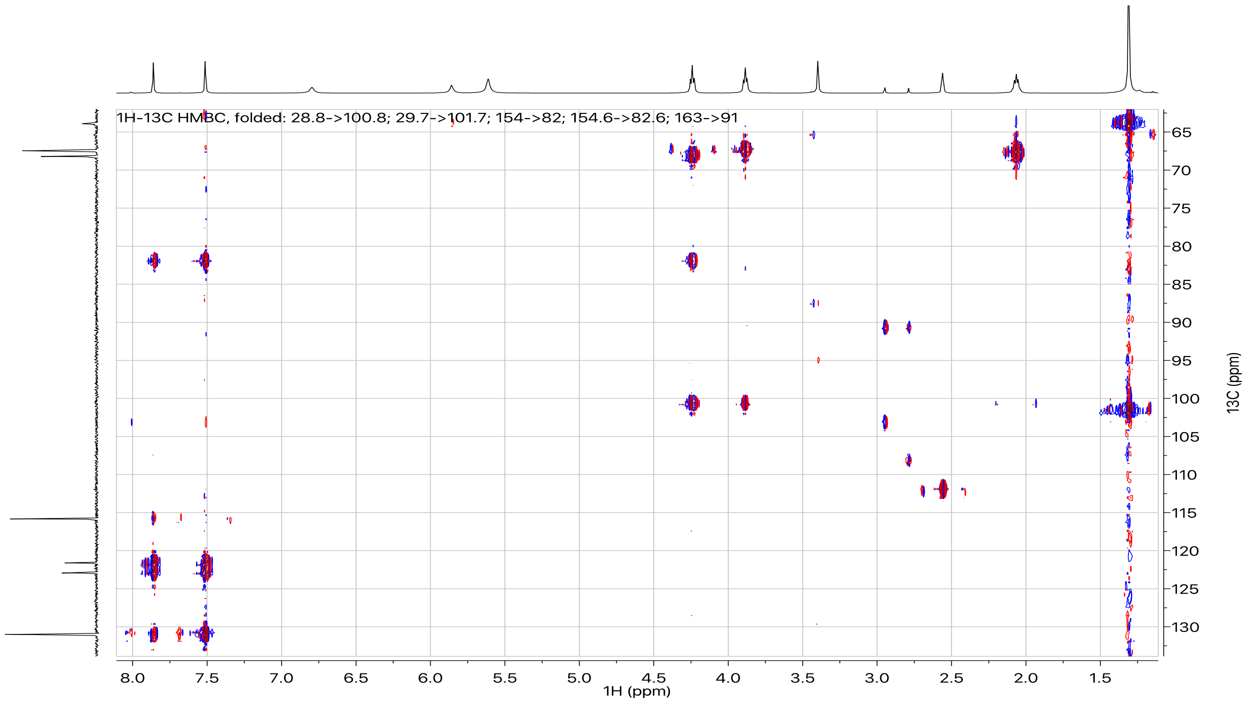

**Figure S18.** $^{1}H$-$^{15}N$ LR-HSQMBC spectrum of **2** in DMSO-$d_6$ at 298K.

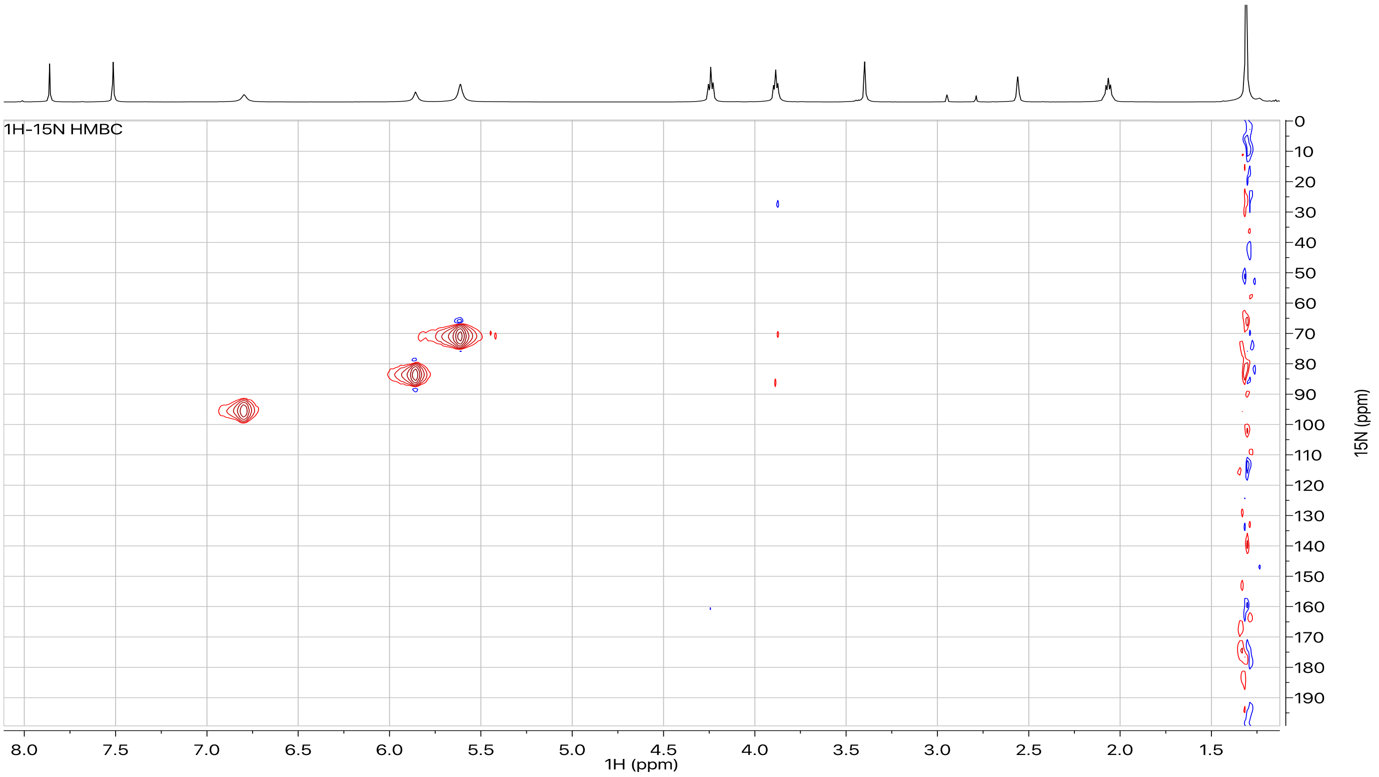

**Figure S19.** HRMS spectrum of **3**.

### Elemental Composition Report

Page 1

#### Single Mass Analysis

Tolerance = 5.0 mDa / DBE: min = -1.5, max = 100.0
Element prediction: Off
Number of isotope peaks used for i-FIT = 3

Monoisotopic Mass, Even Electron Ions
169 formula(e) evaluated with 2 results within limits (up to 50 closest results for each mass)
Elements Used:
C: 0-100 H: 0-250 N: 3-5 O: 0-60 35Cl: 3-3
SG-19DEC19-4-S1-SAM 197 (3.349) AM2 (Ar,25000.0,0.00,0.00); ABS
TOF MS ES+

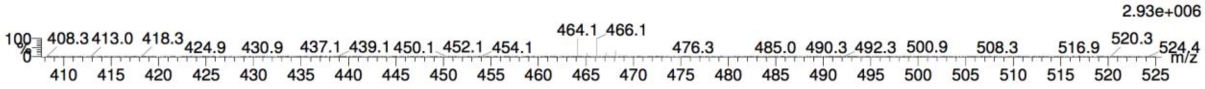


| | | | | | | | | |
|---|---|---|---|---|---|---|---|---|
| Minimum: | | | | -1.5 | | | | |
| Maximum: | | 5.0 | 5.0 | 100.0 | | | | |
| Mass | Calc. Mass | mDa | PPM | DBE | i-FIT | Norm | Conf(%) | Formula |
| 464.1387 | 464.1387 | 0.0 | 0.0 | 6.5 | 630.4 | 0.532 | 58.76 | C19 H29 N5 O2 35Cl3 |
| | 464.1427 | -4.0 | -8.6 | 10.5 | 630.7 | 0.886 | 41.24 | C24 H29 N3 35Cl3 |

**Figure S20.** $^{1}$H NMR spectrum of **3** in DMSO-$d_6$ at 297.9K.

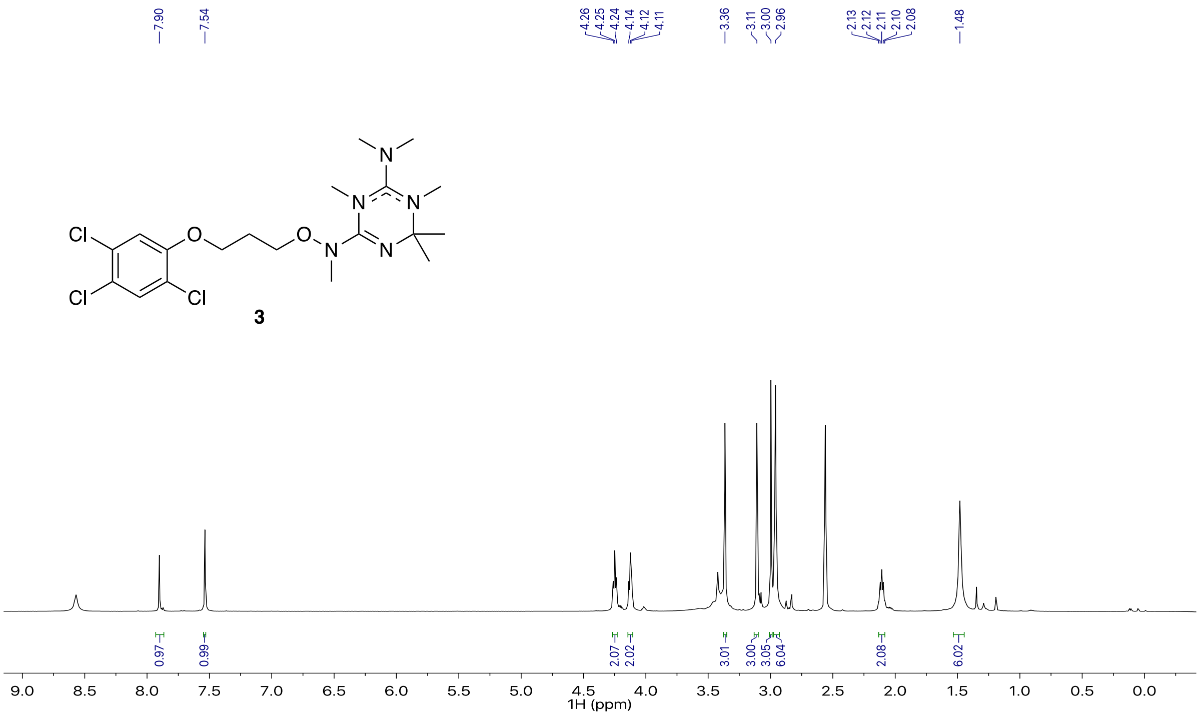

**Figure S21.** $^{13}C$ NMR spectrum of **3** in DMSO-$d_6$ at 298K.

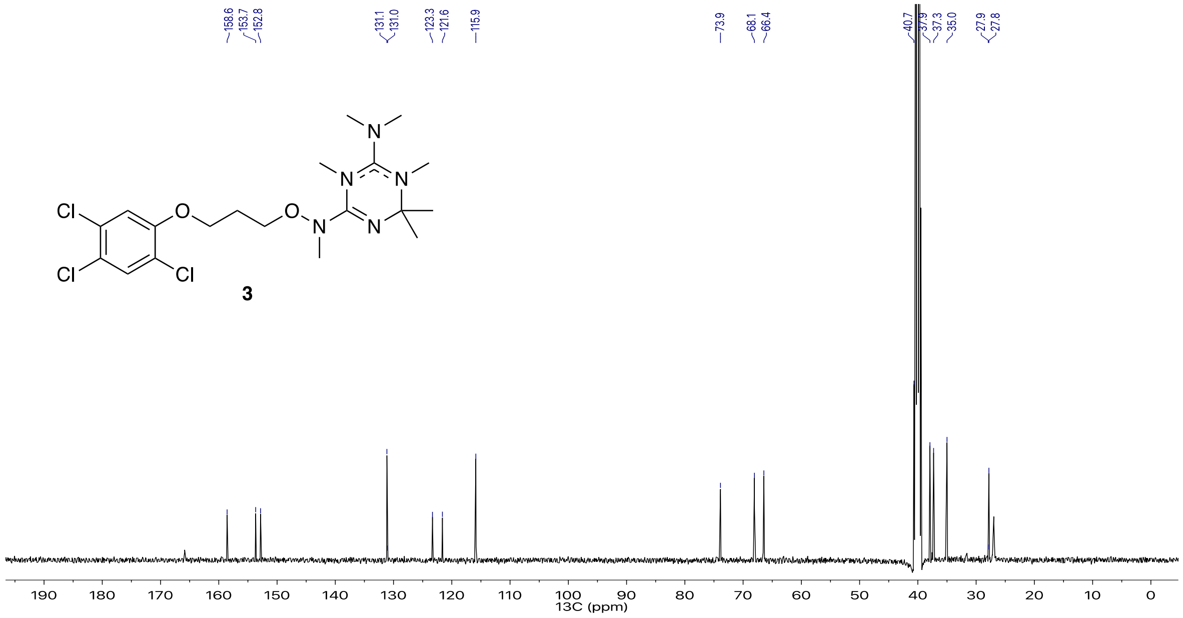

**Figure S22.** $^{1}H$-$^{13}C$ HSQC coupled spectrum of **3** in DMSO-$d_6$ at 298K.

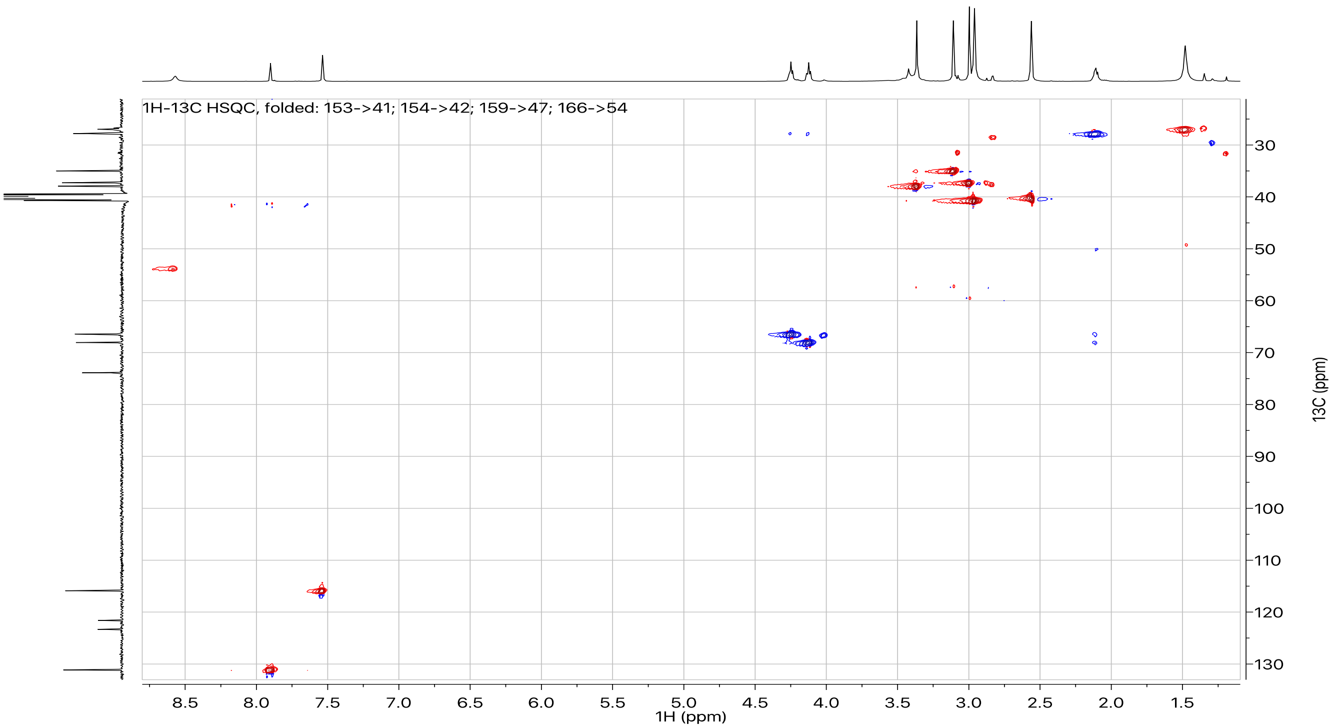

**Figure S23.** $^{1}H$-$^{13}C$ HMBC coupled spectrum of **3** in DMSO-$d_6$ at 298K.

Recorded with a $^{13}C$ sweep width of 72 ppm and the carrier (O2P) set at 99 ppm; folded peaks are labeled on the spectral heading.

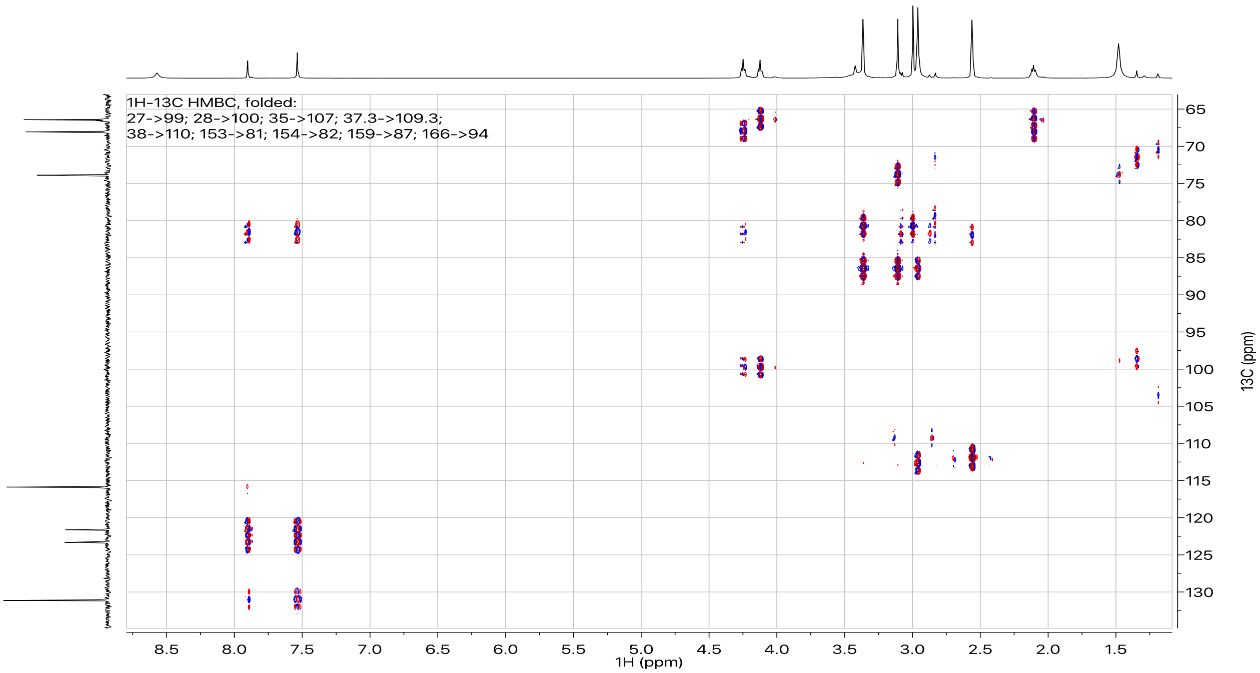

**Figure S24.** $^{1}H$-$^{15}N$ HMBC coupled spectrum of **3** in DMSO-$d_6$ at 298K.

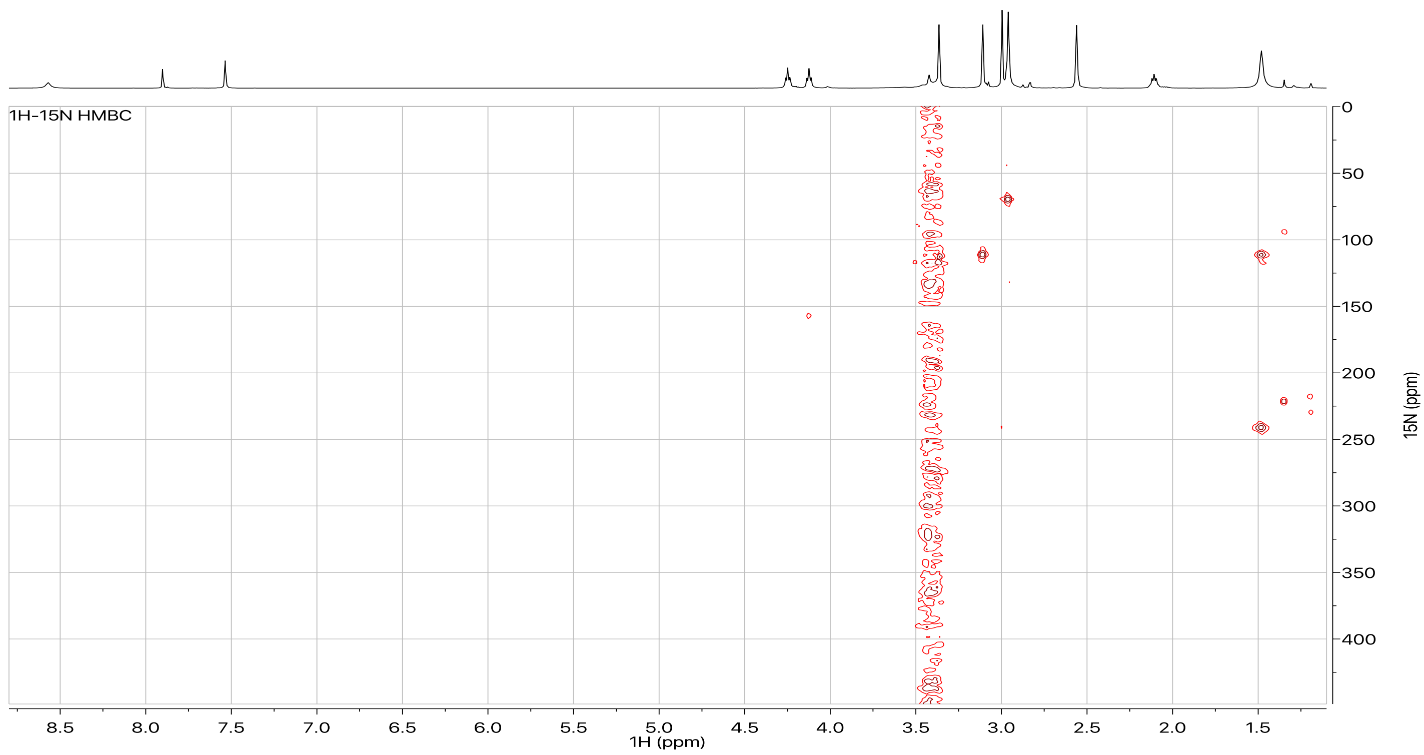

**Figure S25.** 1-D $^1$H NOE spectrum of **3** in DMSO-$d_6$ at 298K. $^1$H's that were irradiated in each spectrum are labeled on the figure.

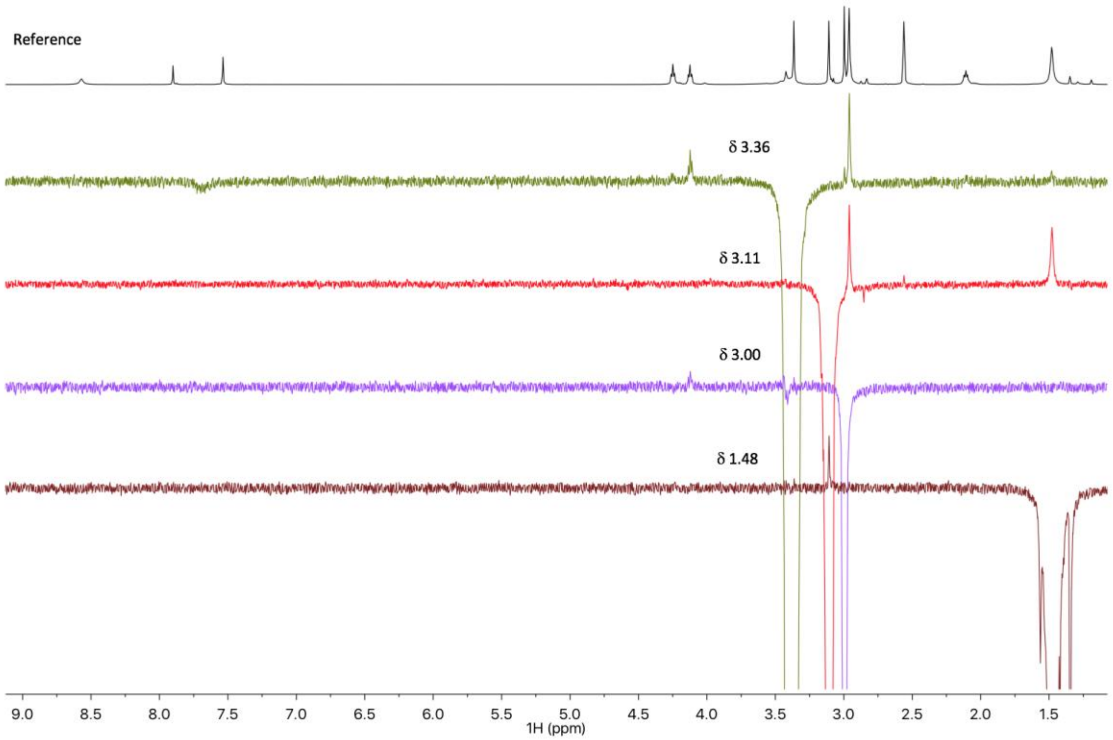
